# A Whole-Genome Approach Discovers Novel Genetic and Non-Genetic Variance Components Modulated by Lifestyle for Cardiovascular Health

**DOI:** 10.1101/700617

**Authors:** Xuan Zhou, Julius van der Werf, Kristin Carson-Chahhoud, Guiyan Ni, John McGrath, Elina Hyppönen, S. Hong Lee

## Abstract

Both genetic and non-genetic factors can predispose individuals to cardiovascular risk. Finding ways to alter these predispositions is important for cardiovascular disease (CVD) prevention. Here, we use a novel whole-genome framework to estimate genetic and non-genetic effects on—hence their predispositions to—cardiovascular risk and determine whether they vary with respect to lifestyle factors. We performed analyses on the Atherosclerosis Risk in Communities Study (ARIC, N=6,896-7,180) and validated findings using the UK Biobank (UKBB, N=14,076-34,538). Cardiovascular risk was measured using 23 traits in the ARIC and eight traits in the UKBB, such as body mass index (BMI), resting heart rate, white blood cell count and blood pressure; and lifestyle factors included information on physical activity, smoking, alcohol consumption and dietary intake. Physical activity altered both genetic and non-genetic effects on heart rate and BMI, genetic effects on HDL cholesterol level, and non-genetic effects on waist-to-hip ratio. Alcohol consumption altered both genetic and non-genetic effects on BMI, while smoking altered non-genetic effects on heart rate, pulse pressure, and white blood cell count. In addition, saturated fat intake modified genetic effects on BMI, and total daily energy intake modified non-genetic effects on waist-to-hip ratio. These results highlight the relevance of lifestyle changes for CVD prevention. We also stratified individuals according to their genetic predispositions and showed notable differences in the effects of lifestyle on cardiovascular risk across stratified groups, implying the need for individualizing lifestyle changes for CVD prevention. Finally, we showed that neglecting lifestyle modulation of genetic and non-genetic effects will on average reduce SNP heritability estimates of cardiovascular traits by a small yet significant amount, primarily owing to overestimation of residual variance. Thus, current SNP heritability estimates for cardiovascular traits, which commonly do not consider modulating effects of lifestyle covariates, are likely underestimated.

## Introduction

Cardiovascular diseases (CVD) are the world’s number one cause of mortality, claiming an estimated total of 17.7 million lives globally in the year 2015 alone—that is 31% of total deaths in just a single year^1^. Managing CVD risk is therefore a top public health priority worldwide. It is estimated that between 20% to 60% phenotypic variability in CVD related traits such as blood pressure and blood clotting factors are due to additive genetic variation (see ^2,3–6^); and the remaining 40% to 80%, commonly referred to as residual variation, could arise from random measurement errors and systematic non-genetic variation in the epigenome, transcriptome, metabolome, proteome and microbiome, which are involved in or interact with the translation of genotype to phenotype.

Given the substantial genetic and non-genetic contributions to CVD risk, identifying ways that modify their effects can have important implications for CVD prevention. In fact, the idea of Genotype-Environment or Genotype-Covariate (G-C) interaction is well established for traits such as BMI^7–9^. That is, genetic effects vary depending on environmental exposure, such as modifiable lifestyle covariates including smoking, alcohol intake and physical activity. Much like G-C interaction to genetic variance, we recently demonstrated that some non-genetic variance component can exist that changes with respect to lifestyle covariates, which we termed Residual-Covariate (R-C) interaction^10^, i.e., phenotypic variation with respect to lifestyle covariates that is independent of genetic effects.

Understanding G-C and R-C interactions in the context of cardiovascular traits will not only translate into empowering public messages but also enable personalized lifestyle changes for CVD prevention according to individuals’ genetic and non-genetic information, as opposed to a one-fits-all approach that neglects individual differences. Aside from its practical implications, studying G-C and R-C interactions is also of theoretical value as it may offer some insight into missing heritability^11,12^.

To date, G-C interaction estimates for cardiovascular traits are based on a limited number of genetic variants^13–20^; therefore they are likely underestimated. R-C interaction has been largely neglected, leading to potential confounding between G-C and R-C interactions, in the presence of genuine R-C interaction^10^. Here, using a novel whole-genome approach^10^, we extend the current understanding of G-C and R-C interactions on cardiovascular health. Instead of focusing on genetic variants with large phenotypic effects, our approach uses all common Single Nucleotide Polymorphisms (SNPs) capturing variation across the entire genome, thereby providing genome-wide estimates of G-C interaction. Further, our approach allows residual variance to vary with respect to a chosen covariate, thereby providing estimates of R-C interaction. By examining G-C and R-C interactions, we aim to identify lifestyle factors that modify genetic and/or non-genetic effects on traits that are indicative of CVD risk.

## Results

### Method Overview

We used Multivariate Reaction Norm Models (MRNMs)^10^ to estimate genetic and residual variance components of cardiovascular traits that vary with respect to lifestyle covariates, which we termed G-C and R-C interactions, respectively (see Methods for details). To detect these two types of interactions, we fitted two MRNMs—one assumes no G-C and R-C interactions (i.e., a null model) and the other assumes both G-C and R-C interactions (i.e., a full model)—and declared the presence of one or more interaction terms when the full model had a better fit than the null (see Supplementary Note 2 for justification). After careful calibration of our model comparison method using simulations, we performed primary analyses on the Atherosclerosis Risk in Communities (ARIC) study, which contains dense cardiovascular health related variables and lifestyle covariates. We chose 23 CVD related traits, including coagulation factors, blood pressure, heart rate, and BMI etc.; and 22 lifestyle covariates that cover physical activity, alcohol intake, cigarette smoking and dietary composition (see Methods for details). Signals emerging from the ARIC dataset were subsequently validated in the UK Biobank (UKBB) dataset, if related data are available.

### Simulation

To calibrate the null versus full model comparison method, we simulated phenotypic data with no G-C and R-C interactions, data with G-C and/or R-C interactions of small and large magnitudes, data that conform to the normality assumption held by MRNMs, and data that do not (see Supplementary Table 1 for details). In each scenario, we repeated the simulation 100 times, resulting in 100 replicates of simulated data. Using these data, we tested the extent to which the true data generating models can be recovered by our model comparison method.

When the model assumption of normality was met, the type I error rate of the null versus full model comparison method was controlled (0.04; see Supplementary Table 2). Small and large phenotypic deviations from the normality inflated the type I error rate to 0.2 and 0.65, respectively. However, after a rank-based inverse normal transformation (RINT) of phenotypic data, the type I error rate was approximately controlled (0.05 and 0.07 for large and small phenotypic deviations from normality, respectively; Supplementary Table 2), indicating that an RINT can effectively reduce false positive findings in face of violations of the normality assumption.

The statistical power of the null versus full model comparison was estimated using data simulated under scenarios other than the null, i.e., G-C only, R-C only and both G-C and R-C interactions. We found that whether the normality assumption is met or not, the proportion of replicates for which the full model had a better fit than the null was at least 0.88 (Supplementary Table 2), giving an estimated power above 88%. Applying an RINT did not affect the power in any scenario.

For each simulation scenario, we compared parameter estimates from the full model with their corresponding true values. Supplementary Figure 1 shows sampling distributions of full-model parameter estimates based on 100 replicates for both large and small effects settings (in terms of heritability, G-C and R-C interactions; Supplementary Table 1) when the model assumption of normality was met, and it indicates that the full model produced unbiased estimates of model parameters under all simulation scenarios. This observation holds even when the normality assumption was violated (Supplementary Figures 2 & 3). In contrast, after applying an RINT, full-model estimates were biased for some model parameters (Supplementary Figures 2 & 3).

In summary, our simulation results indicate that when the model assumption of normality is met, the likelihood ratio test that compares the full model with the null can detect G-C and/or R-C interaction at an acceptable type I error rate with a reasonable level of power. When the normality assumption is violated, however, type I error rate would be inflated, in which case an RINT of the phenotype data is an effective remedy without compromising statistical power. In situations where the normality assumption is not violated, a rank-based inverse normal transformation of the phenotype data would not adversely affect type I error rate and statistical power. In terms of parameter estimates, full-model estimates of heritability, G-C and R-C interactions are unbiased, regardless of whether the normality assumption is violated or not. Full-model estimates would however become biased after an RINT. Therefore, for analysis of real data, if the model assumption of normality is in doubt, rank-based inverse transformation should be applied to control type I error rate; and once a significant finding is declared, full model estimates of parameters from data without the transformation should be reported and interpreted.

### G-C & R-C Interactions

For analysis of real data, we had a total of 23 CVD traits, and for each trait, we screened 22 available lifestyle covariates for G-C and R-C interactions. Out of the 506 pairs of cardiovascular trait and lifestyle covariate, 214 yielded significant results at the 0.05 level, where the full model had a better fit than the null; and after Bonferroni correction 68 pairs remained significant (Figure 1). Of these, 34 survived the sensitivity analysis, where we applied an RINT to all traits. In a further investigation, we noted that a large majority of the signals that were lost after the RINT were from traits that have large skewness and kurtosis (Supplementary Figure 4). Given that RINT can control type I error rate when the normality assumption of MRNM is violated, as shown by simulation results, the lost signals are likely to be spurious. Hence, in the following we will focus on signals remained after the RINT.

**Figure 1.**
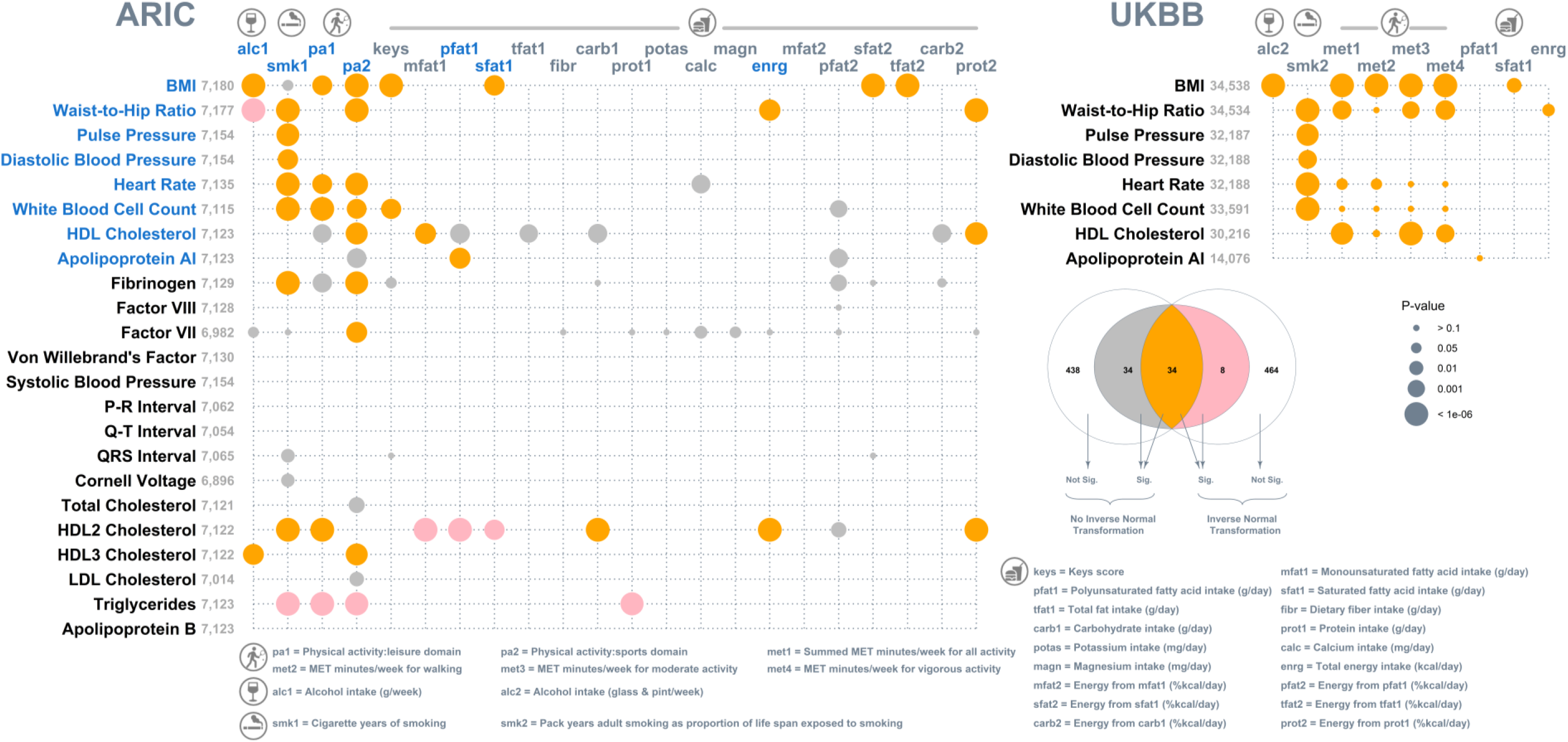
Bubble plot of p values that identify lifestyle modulation of genetic and non-genetic effects on cardiovascular traits. Left: for each of the twenty-three cardiovascular traits (along the y axis) from the ARIC dataset, twenty-two lifestyle covariates (along the x axis) were screened separately for Genotype-Covariate (G-C) and Residual-Covariate (R-C) interactions, by comparing a multivariate reaction norm model that allows G-C and R-C interactions (i.e., a full model) with a null model that assumes no G-C and R-C interactions. The 506 null versus full model comparisons were repeated after a rank-based inverse normal transformation was applied to all traits for a sensitivity analysis. Signals (after Bonferroni correction) for data before and after the transformation are color coded, as detailed in the Venn diagram. Thirty-four signals (in orange) remained after the sensitivity analysis. Seventeen of these remaining signals were subject to validation using the UK biobank, and their corresponding traits and lifestyle covariates are highlighted in blue. Top right: results of the UK biobank validation. For both datasets, bubbles are proportional to p-values based on data after the rank-based inverse normal transformation. Note exceptions to the sample size displayed for BMI vs. sfat1 (N=16,257) and for waist-to-hip ratio vs. enrg (N=16,254) in the UK biobank due to limited availability of dietary intake data among the selected participants.

Out of the 34 significant pairs remaining after the RINT, 17 were covered by the UKBB, allowing replication of the analyses conducted in the ARIC. The majority of these signals, 14 out of 17, were present in both datasets (Figure 1 right). The three signals lost in the replication were the modulating effects of physical activity on white blood cell count and of polyunsaturated fatty acid intake on apolipoprotein a1. In addition, among the replicated signals, results for physical activity varied slightly when METs were broken down into walking, moderate and vigorous activities, indicating that modulating effects of physical activity may be conditional on the type of activity. The variance estimates from the full model for all signals from the ARIC and UKBB datasets are listed in Supplementary Tables 5 and 6, respectively. In summary, our results indicate that lifestyle factors that include alcohol intake, smoking, physical activity, and dietary composition are highly relevant to inter-individual variability in cardiovascular health and hence CVD risk.

For the 34 signals emerged from the ARIC, magnitudes of full-model variance estimates of G-C and R-C interactions were further examined, as proportions of estimated 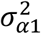 and 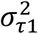 out of total phenotypic variance, respectively, as shown in Figure 2. Four major observations emerged. First, G-C and R-C interactions are sizeable, which can account for up to 20% of phenotypic variance, highlighting the importance of lifestyle modulation to inter-individual variability in cardiovascular health. Second, variance estimates of R-C interactions are in general larger than G-C interactions, indicating that lifestyle covariates play a greater role in modulating non-genetic effects on cardiovascular health than genetic effects. Third, some variance estimates can be zero or even below zero. This is not totally unexpected though and is within the observed range of sampling errors from analyses of simulated data (Supplementary Figure 1). Lastly, we noted a strong inverse correlation between variance estimates of R-C and G-C interactions (Pearson *r*=−0.81). Such collinearity is likely due to the same covariate being used for estimating G-C and R-C interactions. Similar observations were noted in each replicate of simulated data, yet both variance estimates of G-C and R-C interactions were unbiased (Supplementary Figure 1). Thus, despite collinearity between variance estimates, estimation accuracy did not appear to be adversely affected. It is noted that the statistical power to separate G-C and R-C interactions can be low, and parameter estimates from models including only G-C or R-C interaction (referred to as ‘G-C only’ and ‘R-C only’ models) can be biased as shown in simulations (see Supplementary Note 2). Consequently, only the null versus full model comparison was chosen to indicate lifestyle modulation. Nonetheless, we compared nested models, i.e., a G-C only model and a R-C only model, with the full model to assess R-C interaction that is orthogonal to G-C interaction and G-C interaction that is orthogonal to R-C interaction, respectively (Supplementary Table 5 for ARIC & 6 for UKBB).

**Figure 2.**
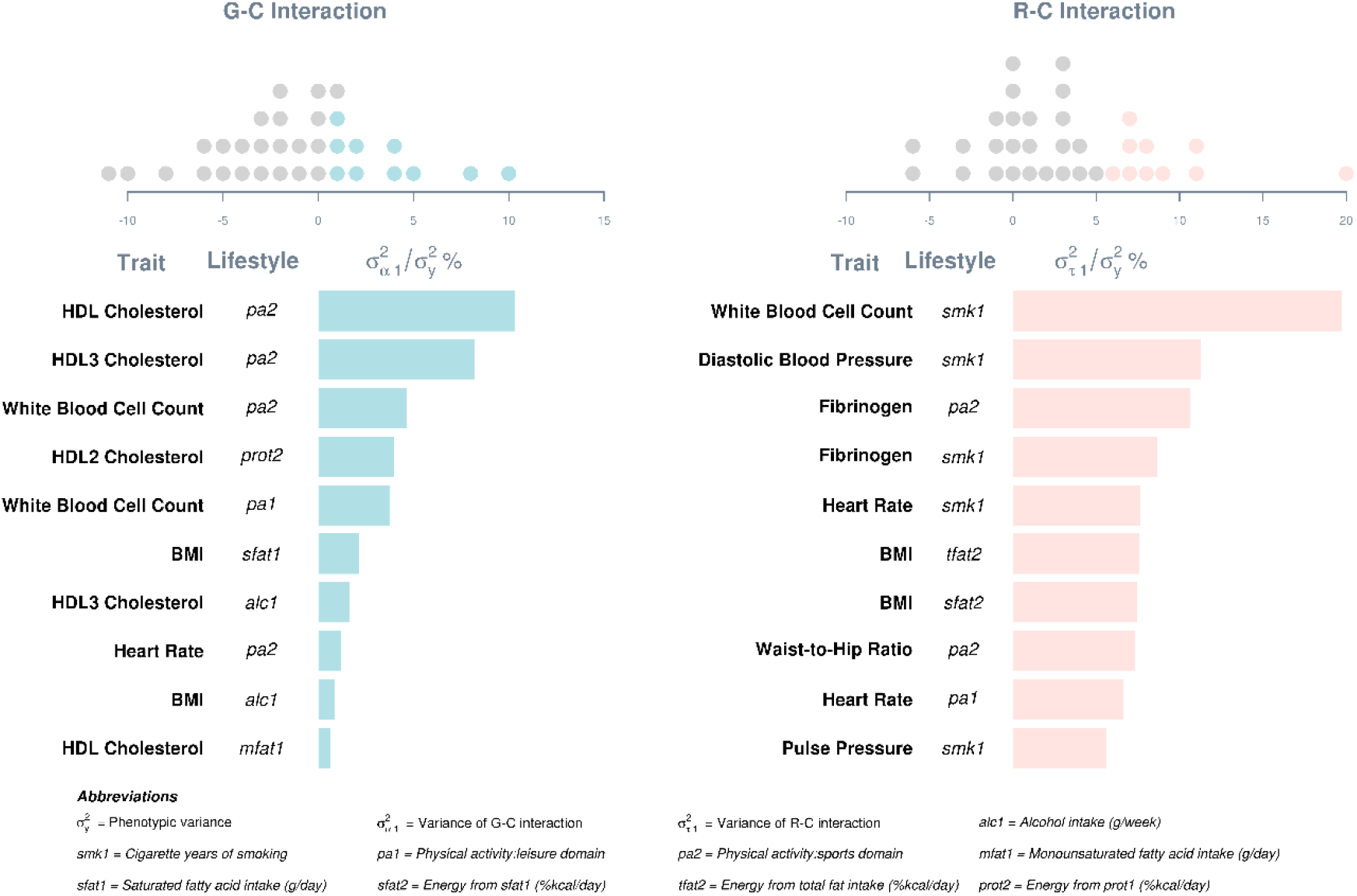
Variance estimates of G-C and R-C interactions as percentages of total phenotypic variance. Estimates were derived by fitting a multivariate reaction norm model that included both G-C and R-C interactions (i.e., a full model) to data without a rank-based inverse normal transformation. Dot plots on the top show distributions of estimates relative to the phenotypic variance of respective traits. Estimates are included only for signals that remained after a sensitivity analysis, where the full model was better than the null after Bonferroni correction on data after a rank-based inverse normal transformation. Top ten estimates are shown in bar charts below.

For the 14 signals that were first discovered in ARIC and replicated in UKBB, we compared variance estimates of G-C and R-C interactions across the two datasets (Supplementary Tables 3 & 4) and noted some similarities. Physical activity altered both genetic and non-genetic effects on heart rate and BMI. It also altered genetic effects on HDL cholesterol level, and non-genetic effects on waist-to-hip ratio. Alcohol consumption altered both genetic and non-genetic effects on BMI, while smoking altered non-genetic effects on heart rate, pulse pressure, and white blood cell count. In addition, saturated fat intake modified genetic effects on BMI, and total daily energy intake modified non-genetic effects on waist-to-hip ratio.

The presence of G-C and R-C interactions indicates heterogeneity of genetic and residual variance-covariance structures with respect to lifestyle covariates^21^, which are depicted in Supplementary Figure 5 for G-C interactions and in Supplementary Figure 6 for R-C interactions. To more explicitly illustrate G-C interactions, for each of the eight traits with the largest variance estimates of G-C interaction, we stratified observations into three groups—top, middle, and bottom—according to per-individual estimate of *α*_1_(via BLUP^22,23^). It is important to note that ***α***_1_ in our model indicates the direction and effect size of G-C interaction for each individual, and it is assumed to follow a normal distribution with mean zero. For each trait, we defined the three groups as having an 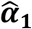 below the 20^th^ percentile (bottom), between the 40^th^ and 60^th^ percentiles (middle), and above the 80^th^ percentile (top), respectively, and plotted their estimated genetic effects, i.e., 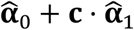, given their standardized values on the relevant lifestyle covariate **c** (Figure 3). Regardless of specific traits, the three groups show distinct trajectories of genetic effects with increasing lifestyle covariate values. Specifically, estimated genetic effects become amplified for the top group, remain stable for the middle group, and become attenuated for the bottom group. Thus, the G-C interactions detected by our models indicate that there exist subpopulations whose genetic effects on cardiovascular traits are modified by lifestyle changes in drastically different ways.

**Figure 3.**
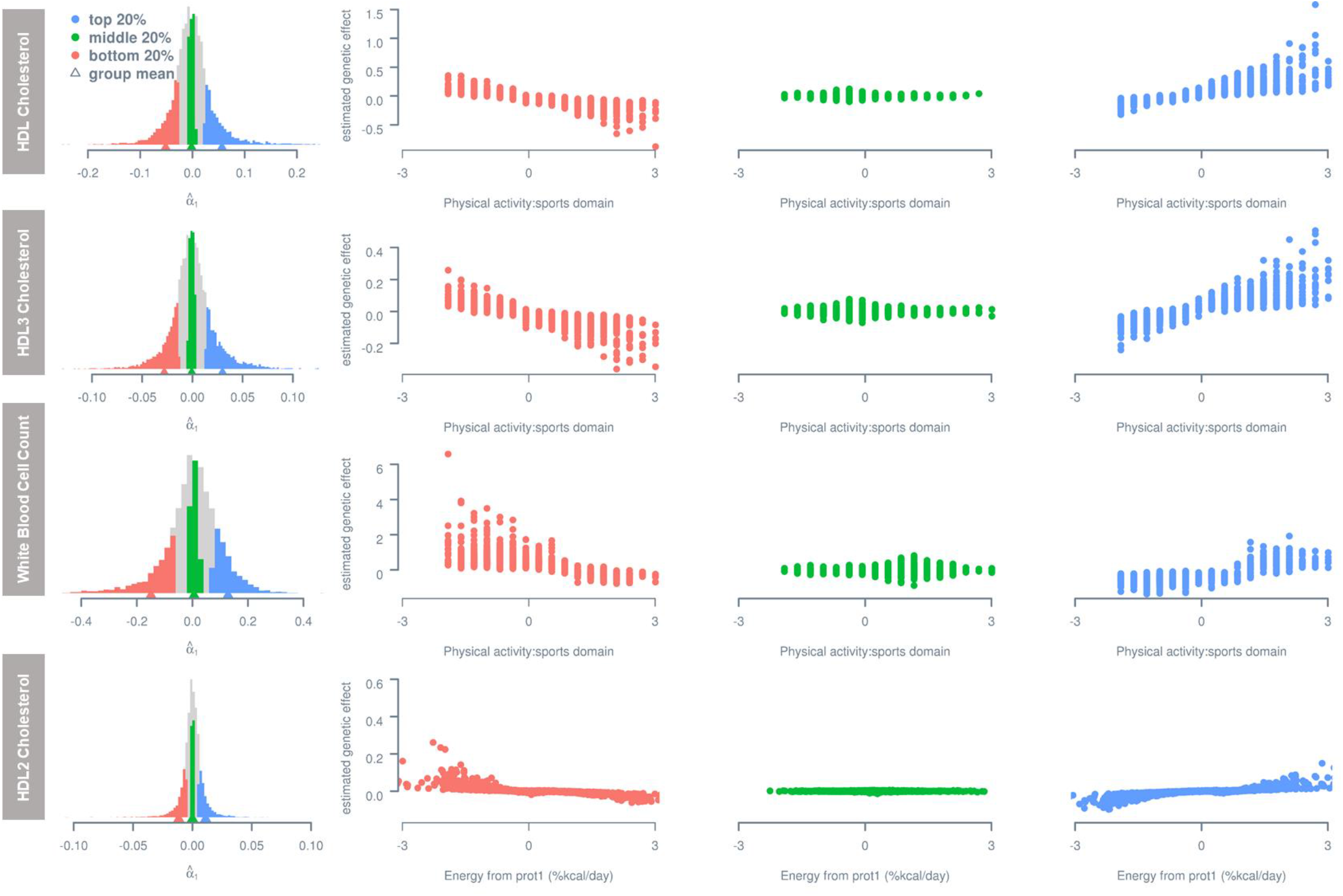
Estimated genetic effects with respect of lifestyle covariate for groups stratified according to per-individual estimate of G-C interaction (i.e., ***α***_**1**_). Histograms on the left show distributions of per-individual estimates of G-C interaction. Only first four traits with the largest variance estimate of G-C interaction are shown. Lifestyle covariates are standardized and are shown in each plot within 3 standard deviations from the mean.

Importantly, the per-individual estimate of *α*_1_that we used for group stratification is an aggregate of G-C interactions over SNPs for the entire genome, hence a genome-wide estimate of G-C interaction. Therefore, individual differences in 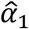 would reflect systematic genetic variation, which would be the most pronounced between the two extreme groups, i.e., top and bottom. A further exploration on pairwise genomic relationship for individuals within and between the two extreme groups revealed that the average genomic relationship within each group is greater than the grand average relationship of the entire dataset, but the average between-group relationship is less than the grand average (Supplementary Table 7). That is, compared to two randomly chosen individuals, a pair of within-group individuals is on average more genetically similar, but a pair of between-group individuals is on average more genetically distant. This observation holds for all eight analyses with the largest variance estimates of G-C interaction (Supplementary Table 7). Thus, the two extreme groups for these analyses in fact have systematic genetic differences.

To explicitly illustrate R-C interactions, for each of the eight traits with the largest variance estimates of R-C interaction, we stratified participants into top, middle and bottom groups according to per-individual estimate of *τ*_1_ in the same way as for G-C interaction. Figure 4 shows estimated residual effects, i.e., 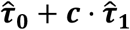, given standardized values on the relevant lifestyle covariate *c* for the three groups. Like for G-C interaction, the three groups show different trajectories with increasing lifestyle covariate values. Thus, as found for genetic variance heterogeneity, residual variance heterogeneity detected as a R-C interaction by MRNMs indicates the presence of subpopulations whose residual effects on cardiovascular traits are modified by lifestyle changes in different ways. Although a genuine R-C interaction can be unbiasedly estimated as shown in the simulation (Supplementary Figure 1), the models used in this study do not inform how individual differences in 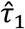 arise, because the fitted variance-covariance structure for residual effects is an identity matrix (see Methods). This problem will no longer exist in a repeated-measures design or if a non-identity matrix is fitted for the variance-covariance structure of residual effects^10^.

**Figure 4.**
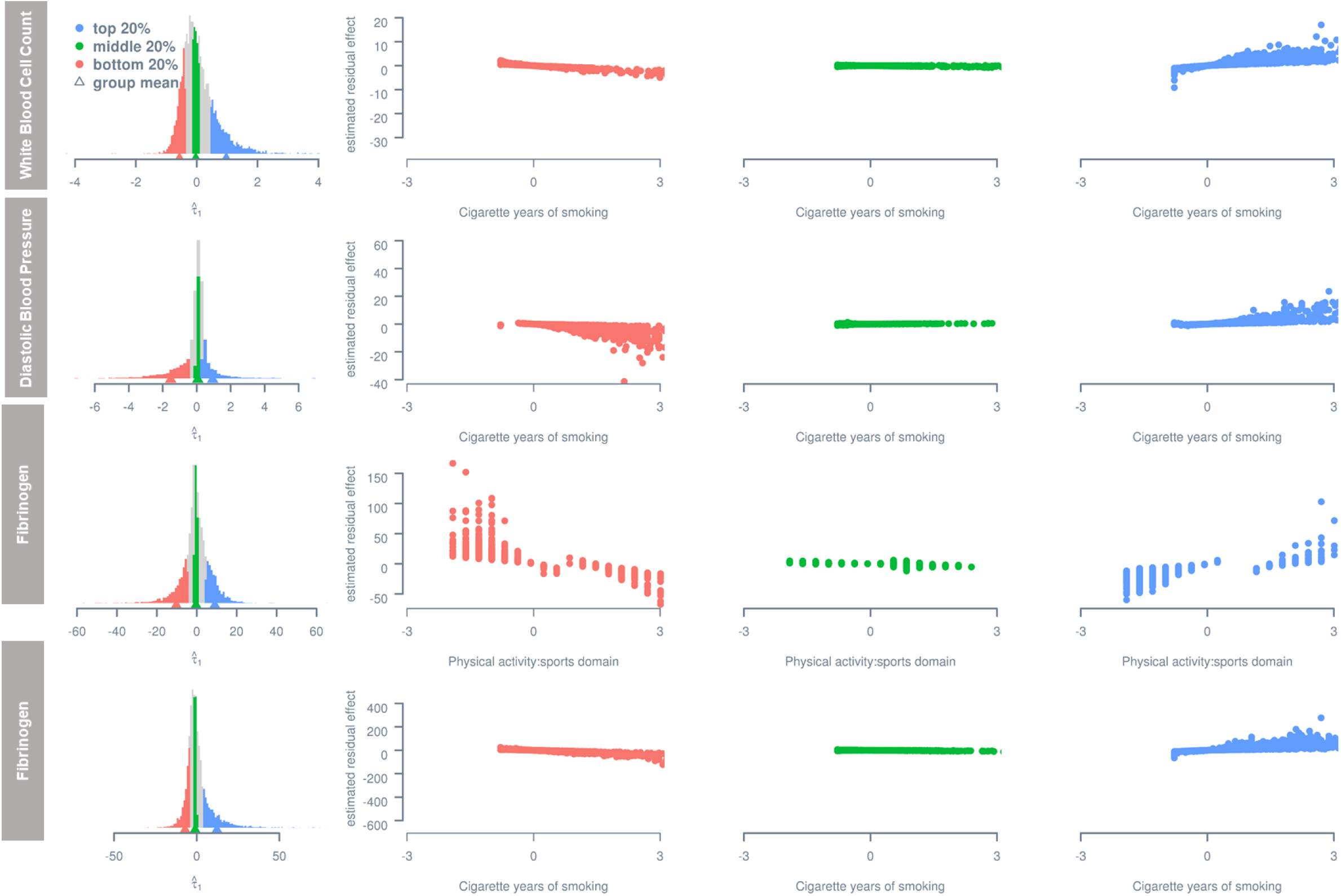
Estimated residual effects with respect to lifestyle covariate for groups stratified according to per-individual estimate of R-C interaction (i.e., ***τ***_**1**_). Histograms on the left show distributions of per-individual estimates of R-C interaction. Only first four traits with the largest variance estimate of R-C interaction are shown. Lifestyle covariates are standardized and are shown in each plot within 3 standard deviations from the mean.

### Heritability

We showed above that lifestyle modulation of genetic and non-genetic effects, in forms of G-C and R-C interactions, is ubiquitous and sizable for cardiovascular traits. To highlight the importance of incorporating lifestyle modulation when estimating trait heritability, we compared SNP heritability estimates from two univariate reaction norm models (URNMs; see Methods for details), one without any interaction terms (i.e., a null model) and the other with one or more interaction terms (i.e., an interaction model) based on results from the multivariate reaction norm models shown above (Figure 1). Null model estimates are essentially equivalent to conventional univariate GREML estimates; hence they are referred to as GREML estimates thereafter. As a contrast, interaction model estimates are thereafter referred to as RNM estimates. Figure 5 is a scatter plot of estimates from both ARIC and UKBB datasets. If GREML and RNM estimates are identical, they are expected to align perfectly along the diagonal line. We found that estimates from the interaction model were, on average, slightly yet systematically larger than estimates from the null model (single-sided paired *t* = 2.35, *df* = 17, *p* = 0.015). Thus, our results support the idea that phenotypic plasticity^21^ can explain some missing heritability (e.g.^24^).

**Figure 5.**
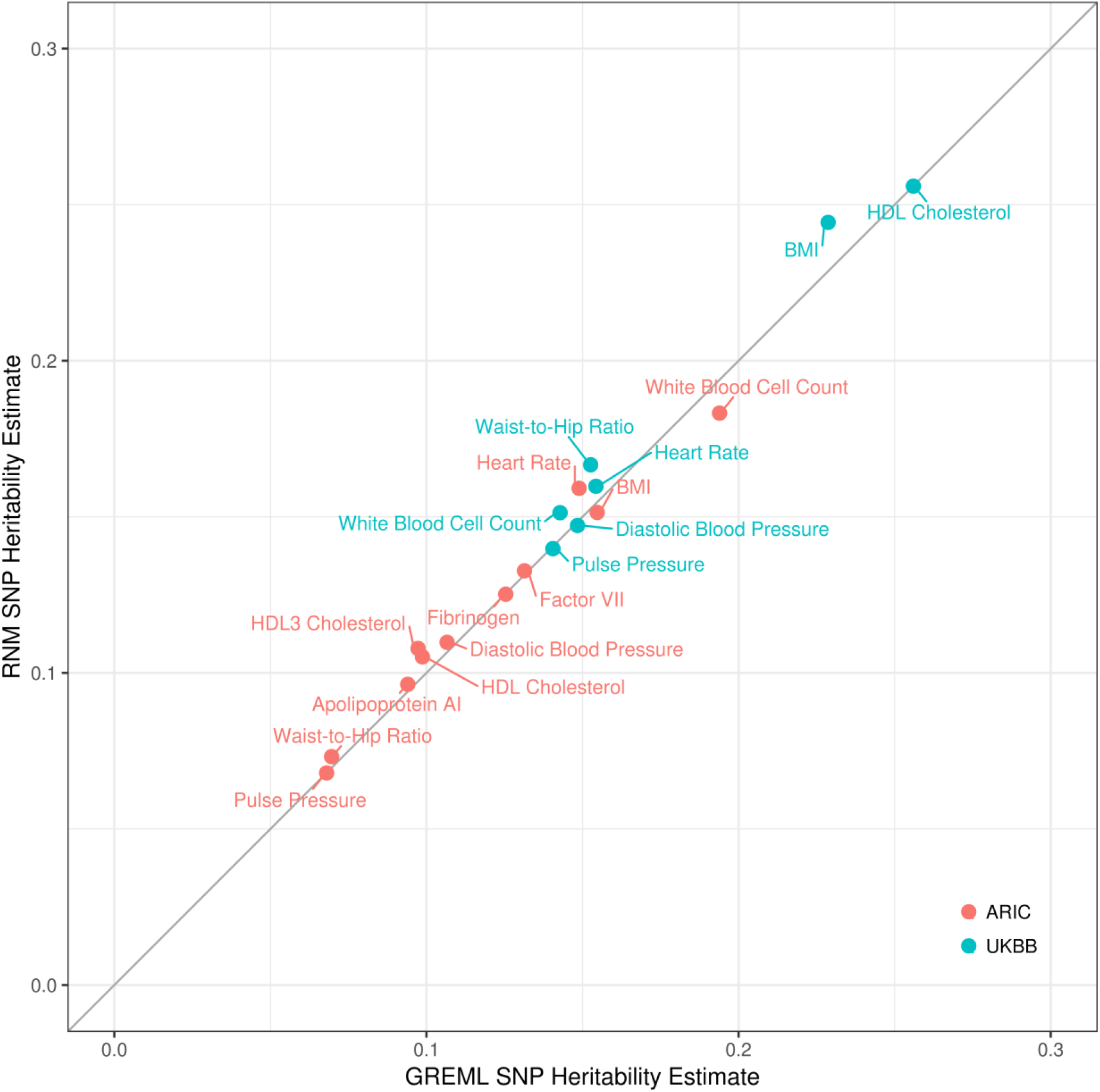
GREML versus RNM SNP heritability estimates. GREML and RNM estimates were derived by fitting a univariate reaction norm model including no interaction term (i.e., null model) and one including one or more interaction terms (i.e., interaction model), respectively. The diagonal is included to highlight the impact of neglecting interaction terms on SNP heritability estimates. Deviations above the diagonal indicate larger RNM estimates relative to GREML estimates. Note HDL2 cholesterol is excluded due to negative variance estimates.

Given that heritability is a function of genetic and residual variance, we further investigated the reason behind larger RNM heritability estimates by comparing GREML and RNM estimates of genetic and residual variance from both ARIC and UKBB datasets (Figure 6). On average, GREML and RNM estimates of genetic variance were not significantly different, but GREML estimates of residual variance were significantly larger than RNM estimates (see mean & 95% CI in Figure 6; two-sided one-sample *t* = 4.15, *df* = 17, *p* = 6.7 x 10^−4^). However, the results from the ARIC and UKBB data sets are somewhat different. In particular, some GREML estimates of genetic variance tend to be underestimated for the ARIC data, which is not evident for the UKBB data. This is likely due to the much smaller sample size of the ARIC data than the UKBB data, which inevitably results in larger sampling errors for the estimates from ARIC. Nonetheless, our results indicate that G-C and R-C interactions are primarily hidden in residual variance estimates in the null model; when they are explicitly estimated in an interaction model, residual variance estimates can be substantially reduced, thereby yielding higher SNP heritability compared to when these components are neglected. This is in line with our previous observation that residual variance is overestimated when fitting a null model to simulated data with genuine G-C or R-C interaction^10^.

**Figure 6.**
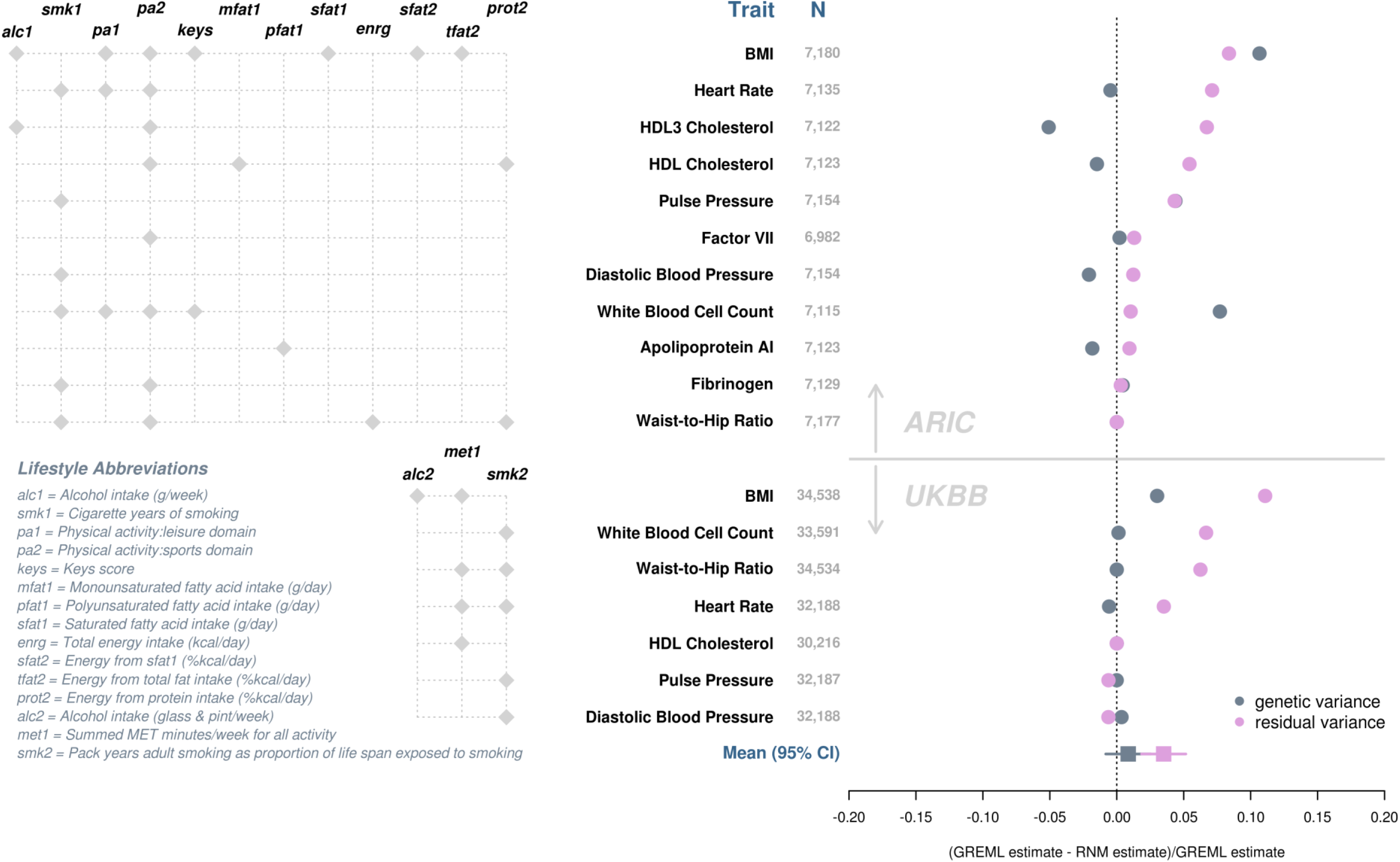
GREML versus RNM estimates of genetic and residual variance. GREML and RNM estimates were derived by fitting a univariate reaction norm model that included no interaction term (i.e., null model) and one that included one or more interaction terms (i.e., interaction model), respectively. The left panel specifies lifestyle covariate(s) included in the interaction model. Changes in genetic and residual variance estimates from the interaction model (i.e., RNM estimates) relative to their respective estimates from the null model (i.e., GREML estimates) are shown on the right to highlight the impact of neglecting interaction terms. Deviations below zero indicate underestimation by GREML, whereas deviations above zero indicate overestimation by GREML. Traits are presented in the decreasing order of deviations for residual variance. Note changes in genetic variance estimates for pulse pressure and waist-to-hip ratio in ARIC and for HDL cholesterol in UK biobank are obscured in the plot by data points for residual variance. HDL2 cholesterol in ARIC was excluded due to negative heritability estimates.

## Discussion

In this study, we used a novel linear mixed model to detect and estimate components of genetic and non-genetic variance that change with respect to modifiable lifestyle covariates, termed as G-C and R-C interactions, in the context of cardiovascular health. Using simulations, we showed that for a sample size of ~7,500 observations, our method has sufficient statistical power to detect genuine G-C and R-C interactions, while keeping the false positive rate controlled. Applying our method to real data, for each of 23 cardiovascular traits selected from the ARIC dataset, we screened for G-C and R-C interactions using 22 available lifestyle covariates that covered smoking, alcohol intake, physical activity and dietary composition. G-C and R-C interactions were found to be ubiquitous among cardiovascular health related traits, and for some traits, estimates were relatively large, accounting for up to 20% of total phenotypic variance.

Among the 14 signals replicated in the UK Biobank, physical activity was found to alter both genetic and non-genetic effects on heart rate and BMI; genetic effects on HDL cholesterol level and non-genetic effects on waist-to-hip ratio. Alcohol consumption altered both genetic and non-genetic effects on BMI, while smoking altered non-genetic effects on heart rate, pulse pressure, and white blood cell count. In addition, saturated fat intake modified genetic effects on BMI, and total daily energy intake modified non-genetic effects on waist-to-hip ratio. To explicitly illustrate G-C and R-C interactions, we stratified individuals according to per-individual estimate of G-C and R-C interactions and showed that genetic and residual effects could take on different directions across groups. While we did not identify any literature in the context of cardiovascular traits that examined R-C interaction, the evidence of G-C interaction in our studies is consistent with the previous literature (e.g.^7,9,13,16–20^), although our study is novel in that G-C interactions were estimated using common SNPs of the entire genome, which are in contrast to estimates based on a single or a limited number of SNPs with large phenotypic effects in past studies.

Given the prevalence of lifestyle modulating effects, we also examined any potential consequence of neglecting these effects on SNP heritability estimates. Such negligence reduced SNP heritability estimates by a small yet significant amount. This reduction is primarily due to overestimation of residual variance. Yet genetic variance estimates are relatively robust to the negligence of significant lifestyle modulation. Our results suggest that current SNP heritability estimates for cardiovascular health related outcomes, which commonly do not take into account modulating effects of lifestyle covariates, are likely underestimated.

Currently, several other approaches to G-C interaction exist in the literature, and our approach is unique in several important ways. Compared to a fixed-effects model approach (e.g.^25^), a mixed-model approach like ours could account for genetic covariance among individuals. Compared to StructLMM^8^, which is a linear mixed-model approach that examines G-C interaction for one SNP at a time, our approach estimates the G-C interaction aggregated over SNPs for the entire genome, thereby providing genome-wide estimates of of G-C interaction. The whole-genome approach to G-C interaction also sets this study apart from G-C interaction studies using a candidate gene approach which only focused on genetic variants with large phenotypic effects (e.g.^13–15^). In these studies, variants might be missed if they contribute to GxE interaction and their effects depend on lifestyle factors. In addition, our approach extends other whole-genome approaches^7,26,27^ by allowing continuous, as opposed to categorical, lifestyle covariates to be modelled; and by simultaneously modelling G-C and R-C interactions.

The prevalence of sizable G-C and R-C interaction effects shown in our study not only reinforces the relevance of existing lifestyle-focused prevention programs for CVD prevention but also suggests that promoting lifestyle changes in a single direction may be ineffective or even inappropriate for some subpopulations. Instead, to most effectively reduce genetic and non-genetic predispositions to unfavourable cardiovascular phenotypes, lifestyle-focused interventions should be tailored to the individual on the basis of his or her relevant genetic and non-genetic information, supporting the rise of precision medicine in cardiovascular disease to individualise treatments and preventions rather than assuming all individuals share a common pathophenotype^28,29^.

Of note, the variance-covariate structure fitted for the genetic effect in our MRNMs is a non-identity matrix constructed using genetic information, i.e., a genomic relationship matrix. In effect, SNP BLUPs derived from MRNMs can be used to predict how a person’s genetic risk would change with respect to a chosen lifestyle covariate, given his or her genetic information. In contrast, the variance-covariance structure fitted for the residual effect in our MRNMs is an identity matrix. Consequently, R-C interactions estimated by our models have little use in the prediction of phenotypes. Further development of MRNMs that incorporate a relationship matrix based on factors underlying residual variations, i.e., a non-identity matrix analogous to a genomic relationship matrix, would be useful for the prediction, and it is currently under way.

As for other approaches to G-C interaction for observational studies, modulating effects of lifestyle covariates found in this study do not imply causality. While randomized controlled trials are the gold standard, further studies using genetic methods such as Mendelian randomization can help determine causal influences. Further, the MRNMs used in this paper are a specific case of the more general MRNMs (see^10^), where genetic and residual effects are expanded to the first order of the chosen lifestyle covariate. Higher order expansions may be necessary and could be employed in future studies where the variance-covariance structure for residual effects is a non-identity matrix. However, increasing model complexity also increases notably the sample size requirement for robust estimation of model parameters. It should also be noted that the sample size in ARIC for our primary analyses is relatively small (6,896–7,180 participants), leading to less precise parameter estimation compared to the UKBB validation analyses. This may explain some discrepancies observed in model estimates between the two datasets. Smoking, for example, was shown to modulate cardiovascular health in both datasets, but the modulations manifested primarily as R-C interactions in ARIC analyses but as G-C interactions in UKBB analyses (see Supplementary Tables 3 & 4). Independent datasets are required to determine the nature of the modulation effects of smoking on cardiovascular traits. Finally, in this paper we only considered intermediate cardiovascular traits that are continuous in nature. Development of valid MRNMs for binary outcome is currently under way. Future applications of these MRNMs would help identify modulating lifestyle covariates that are directly relevant to cardiovascular disease outcomes.

In summary, we found strong modulations from lifestyle covariates, including smoking, alcohol intake, physical activity and dietary composition, for genetic and residual effects on phenotypes that are known to associate with cardiovascular diseases. To illustrate these interactions, we showed that genetic and residuals effects—which may be interpreted as genetic and non-genetic predisposition to CVD health risk, respectively—could change with respect to lifestyle change in different directions for different individuals. Our findings, therefore, reinforce the relevance of lifestyle changes to cardiovascular health and highlight the need for individual considerations when designing lifestyle intervention programs to effectively reduce genetic and non-genetic predisposition to unfavourable cardiovascular phenotypes. Future investigations into specific genetic and non-genetic factors that give arise to individual differences in CVD health risk trajectories with respect to lifestyle changes are well warranted.

## Methods

Our analyses were based on two datasets, the Atherosclerosis Risk in Communities (ARIC) study and the UK Biobank (UKBB). The former was chosen for our primary analyses, because it covers a wider range of cardiovascular traits than the latter. The UKBB, which has a larger sample size than the ARIC study, was chosen as the validation set. Sample size for our analyses depended on availability of phenotype and genotype data, which varied between studies and cross traits. For ARIC, sample sizes were between 6,896 and 7,180. For UKBB, sample sizes were between 14,076 and 34,538.

### ARIC

The ARIC study is a prospective study on the aetiology of atherosclerosis, with data collected from up to five visits over fifteen years from participants of four U.S. communities (Forsyth County, North Carolina; Jackson, Mississippi; suburbs of Minneapolis, Minnesota; and Washington County, Maryland), who were aged between 45 and 64 years in 1987-89^30^. To maximize the sample size for our analyses, we only used data from visit one that occurred in 1987-89. The following phenotypic traits were selected for analyses: plasma concentration/activity of four coagulation factors, namely factor VIII, factor VII, Von Willebrand's factor and fibrinogen; four electrocardiography-derived variables, namely P-R interval, Q-T interval, QRS interval, and Cornell voltage; three blood pressure measures, namely systolic and diastolic blood pressure and pulse pressure; resting heart rate, total cholesterol and total triglycerides levels and white blood cell count. In addition, we also included two anthropomorphic measures: body mass index (BMI) and waist-to-hip ratio. All available lifestyle covariates that had sufficient data were chosen for our analyses. This included smoking, alcohol intake, dietary composition, and physical activity. Detailed descriptions of the selected traits and lifestyle factors are listed below.

#### Cardiovascular Traits

A total of 23 cardiovascular health related traits were selected. Coagulation factors were determined in the ARIC Central Hemostasis Laboratory using previously published procedures^31^. Plasma concentration of fibrinogen was measured by the thrombin-titration method, factor VII and factor VIII activity by clotting assays, and Von Willebrand’s factor antigen with an ELISA technique ^32,33^. P-R interval, Q-T interval, QRS interval, and Cornell voltage were derived from standard 12-lead electrocardiography^34,35^. Sitting blood pressure (systolic and diastolic) was measured three times from the right arm and calculated based on the average of the last two readings. Pulse pressure was computed as the difference between systolic and diastolic blood pressure.

#### Lifestyle Covariates

A total of 22 lifestyle covariates were selected. Smoking was indexed by ‘cigarettes per year’, derived by multiplying the average number of cigarettes per day with the number of years smoked. Alcohol intake was indexed by usual ethanol intake from wine, beer and hard liquor in grams per week. Diet was assessed using a 66-item food-frequency questionnaire based on the Willett 61-item questionnaire^36^. Summary measures derived included dietary lipid content, as indexed by the keys score^37,38^, daily dietary intake of fibre, monounsaturated, polyunsaturated, and saturated fatty acids, total fat, carbohydrate, protein, potassium, calcium, and magnesium; total daily energy intake; and percentages of daily total energy intake from monounsaturated, polyunsaturated, and saturated fatty acids, total fat, carbohydrate, and protein. Physical activity was assessed in work, sports and leisure domains using a modified Baecke questionnaire^39,40^. Only summary scores from sports and leisure questions were used. The score for sports is a summary of 1) the frequency, duration, and an assigned intensity of the sports reported by participants and 2) three additional questions on frequency of sweating, general frequency of playing sports, and a self-rating of the amount of leisure time physical activity compared with others of the same age[s]^41^. The score for leisure is a summary of frequency of watching television (scored inversely), walking, bicycling, and walking/biking to work or shopping^41^.

#### Genotyping Data

The ARIC genotype dataset contains 609,441 single nucleotide polymorphisms (SNPs) from 8,291 participants. We first selected autosomes from white participants then applied standard quality control procedures to the selected dataset. This involved 1) excluding SNPs that do not have a location; ones that have a genotyping rate less than 95%; ones that failed the Hardy-Weinberg test at the 0.0001 level; and ones that had a frequency less than 0.01; 2) excluding individuals who were missing 5% of genotype data; and 3) removing related individuals by excluding one person, at random using a Bernoulli distribution with a selection probability of 0.5, from each pair that had an estimated genomic relationship^26^ greater than 0.05. Eventually, 586,257 SNPs from 7,513 individuals remained. Among these individuals, 6,896 to 7,180 have phenotype data available for analysis.

### UKBB

The UKBB contains health-related data from ~ 500,000 participants aged between 40 and 69, who were recruited throughout the UK between 2006 and 2010^42^. For validation purposes, we only selected phenotypes and lifestyle covariates that overlap with the ARIC dataset, which included BMI, waist-to-hip ratio, heart rate, white blood cell count, diastolic and systolic blood pressure, pulse pressure, HDL cholesterol level, apolipoprotein a1 level, smoking (pack years of smoking as proportion of life span exposed to smoking), alcohol intake (average weekly intake of all types) and physical activity (metabolic equivalent or MET minutes for walking, moderate activity, vigorous activity, and all types^43^), and dietary composition (polyunsaturated fatty acid, saturated fatty acid, and total energy intake).

#### Genotyping Data

The second release of the UKBB genotyping dataset was used. Before quality control, the data set contains 92,693,895 imputed autosomal SNPs from 488,377 individuals. We selected HapMap3 SNPs from individuals of white British ancestry only and applied the same quality control procedures as for the ARIC genotyping dataset (see above). Only HapMap3 SNPs were selected because they were shown to yield reliable and robust estimates of SNP-based heritability and genetic correlation^44–46^. In addition, ambiguous and duplicated SNPs and SNPs with an information score (used to index the quality of genotype imputation) less than 0.6 were excluded. We computed the genomic relationship matrix of all observations and excluded population outliers, defined as individuals who have a score outside three standard deviations on either the first or second principal component of the genomic relationship matrix. From remaining participants, only those who were part of the first release of the UKBB genotyping data (~150,000 individuals) were selected for the purpose of reducing computational burden. This subset of participants therefore has two versions of imputed genotyping records, one from each release, which enabled computation of discordance rates between the two versions for each SNP across individuals and for each individual across SNPs. SNPs and individuals who have a discordance rate > 0.05 were excluded. Eventually, 1,130,918 SNPs from 66,281 participants remained. Among these participants, only 14,076 to 34,538 have phenotype data available for analysis.

### Statistical Analyses

#### Statistical Models

We used a novel whole-genome modelling framework, Multivariate Reaction Norm Models (MRNMs^10^), to detect G-C and R-C interactions. MRNM is an extension of bivariate linear mixed models. In the simplest form of a bivariate linear mixed model, the main trait, *y*, and the covariate, *c*, for individual *i*, after adjusting for their respective fixed effects, *μ*_*y*_ and *μ*_*c*_, are simultaneously expressed as

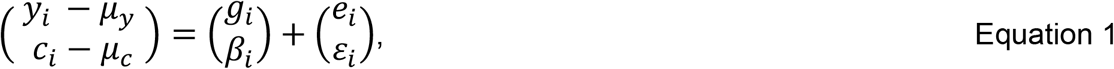

Where 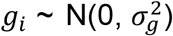 and 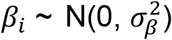 are genetic effects, which are aggregates of random effects of genome-wide SNPs on the main trait and on the covariate, respectively; 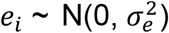 and 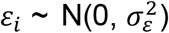 are residual effects, and both *g*_*i*_ and *β*_*i*_ are independent from *e*_*i*_ and *ε*_*i*_.

MRNM extends Equation 1 by introducing random regression coefficients of the covariate, which can be written as

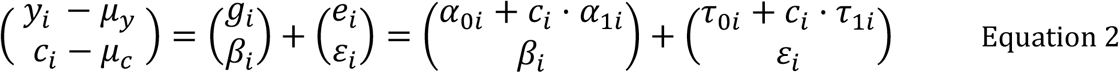

Where *g*_*i*_ breaks into *α*_0*i*_ + *C*_*i*_ · *α*_1*i*_, *e*_*i*_ into 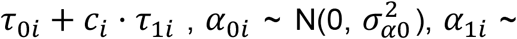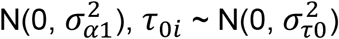 and 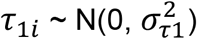.

As shown in both equations, variance of the main trait and of the covariate are partitioned into two general sources, one of genetics (i.e.,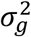 & 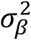) and one of non-genetics or residuals (i.e.,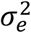 & 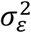). By modelling the main trait and the covariate simultaneously, the covariance between the main trait and the covariate, in forms of *cov*(*g*_*i*_, *β*_*i*_) and *cov*(*e*_*i*_, *ε*_*i*_), is accounted for in a MRNM^10^. This is important given that the covariance between the main trait and covariate can sometimes be nontrivial and would have been neglected in univariate random regression models. More importantly though, the *g*_*i*_ and *e*_*i*_ terms are expanded in terms of *c*_*i*_ in Equation 2, which offers opportunities to model the genetic and residual variances of the main trait as a function of the covariate. With this expansion, it is immediately clear that genetic variance 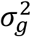 breaks into var(**α**_0_ + *c* · **α**_1_) and residual variance 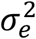 into var(*τ*_0_ + *c* · *τ*_1_), both of which vary with respect to the covariate. As such, MRNMs can estimate and detect genetic and residual variance heterogeneity due to the chosen covariate. A G-C interaction that underlies genetic variance heterogeneity is indicated by significant 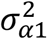 and cov(*α*_0_, *α*_1_), and a R-C interaction that underlies residual variance heterogeneity is indicated by significant 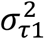 and cov (**τ**_0_, **τ**_1_).

The MRNM shown in Equation 2 is referred to as the **full** model, which assumes and detects both genetic and residual variance heterogeneity with respect to the covariate. The full model can be simplified into other three major forms. Specifically, by setting both var(*c* · *α*_1_) and var(*c* · *τ*_1_) to 0, the **null** model assumes no heterogeneity in either the genetic or the residual variance of the main trait with respect to the covariate. By setting var(*c* · *τ*_1_) to 0, the **G-C** model assumes no R-C interaction and estimates the extent of genetic heterogeneity with respect to the covariate. Finally, by setting var(*c* · *α*_1_) to 0, the **R-C** model assumes no G-C interaction and estimates the extent of residual heterogeneity with respect to the covariate. Detailed descriptions on the variance-covariance structure assumed by MRNMs are included in Supplementary Notes and can also be found elsewhere ^10^.

Prior to model fitting, we attempted to simplify the general MRNMs outlined above by reducing the number of free parameters for estimation. We estimated heritability of each lifestyle covariate in the ARIC dataset via univariate Genomic Restricted Maximum Likelihood (GREML), and found that all estimates were close to zero. Daily potassium intake was the only covariate with an estimate marginally different from zero (*h*^2^ = 0.08 ± 0.04). Subsequently, we simplified MRNMs by setting 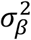 (i.e., genetic variance of the covariate) and its associated covariance terms, i.e., *cov*(*α*_0_, *β*) and *cov* (*α*_1_, *β*), to 0. Unless specified otherwise, all MRNMs fitted to ARIC data in this paper are simplified MRNMs.

For each pair of main trait and covariate, the null and full models were fitted and compared using a likelihood ratio test. For the simplified MRNMs, the test statistic, i.e., −2 log likelihood ratio, is assumed to have a chi-square distribution with five degrees of freedom. The alpha level was set at 0.05. A significant p-value indicates the full model has a better fit than the null, hence the presence of a G-C, R-C interaction, or both. Since the full model does not separate G-C and R-C interactions, we considered a model comparison strategy to separate the two. However, we show in the Supplementary Notes that this strategy can suffer from weak statistical power and biased estimation, which makes it an overall inferior method to the null versus full model comparison method for detecting G-C and R-C interactions. Therefore, our results are based on the latter. All model fitting for this paper was performed using MTG2 ^47^.

#### Adjustments for Main Traits & Lifestyle Covariates

Before fitting MRNMs, all main traits were adjusted for demographics and lifestyle factors using fixed-effects linear models. In effect, changes in the variance-covariance structure of the main trait with respect to a given covariate were not confounded by adjustment factors or by any unobserved variables that linearly covary with adjustment factors. The demographic variables for main trait adjustments included age, sex, education level, marital status, field centre ID, and population structure, as measured using the first fifteen principal components of the estimated genomic relationship matrix. All lifestyle factors described in the previous section were used for main trait adjustment. Some additional adjustment factors were also included, depending on the main trait. For heart rate, blood pressure measures, electrocardiography variables and coagulation factors, additional adjustment factors included hypertension, defined as systolic blood pressure >=140 or diastolic blood pressure >= 90, and hypertension lowering medication use. For total cholesterol and triglycerides levels, additional adjustment factors were hypertension, hypertension lowering medication use, cholesterol-lowering medication within two weeks, and medications that secondarily affect cholesterol. As the second trait in the multivariate reaction normal model (see the second part of Equation 1), lifestyle covariates were pre-adjusted in the same way as the main trait in the first part of Equation 1.

### Heritability

We considered the consequence of neglecting G-C and R-C interactions on heritability estimates. Specifically, we estimated heritability of each trait using two models, one that includes no interaction term at all, i.e., null model (also known as GREML), and the other that includes one or more interaction terms, i.e., interaction model, and compared estimates of the two models. To reduce computational burden, we used univariate reaction norm models (URNMs), as opposed to MRNMs. The null model in the univariate framework is essentially Equation 1 without the part that involves the covariate, *c*_*i*_. Using the same notation as Equation 1, the main trait for individual *i*, in a URNM can be written as:

Null model:

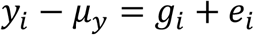

The interaction model in the univariate framework expands *g*_*i*_ and *e*_*i*_ as functions of *m1* and *m2* covariates, respectively, where *m*_1_ + *m*_2_ ≥ 1. Using *j* to index covariate, the main trait for individual *i* in a URNM with interaction terms can be written as:

Interaction model:

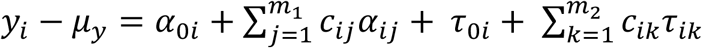

### Simulation

To facilitate data interpretation, we simulated phenotypic data with and without G-C and/or R-C interactions and assessed, using simulated data, whether MRNMs can produce unbiased parameter estimates, type I error rate and power of detecting G-C and R-C interactions. We purposely chose two sets of model parameter configurations that varied primarily in effect size for heritability, G-C and R-C interactions. One setting had large effect sizes, referred to as the ‘large-effects setting’, with a heritability of 0.5 for both the main trait and the covariate and both 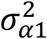 and 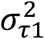, which are indicative of G-C and R-C interactions, were set at 0.5. In contrast, the other setting had smaller effect sizes, referred to as the ‘small-effects setting’, with a heritability of 0.15 for the main trait, 0 for the covariate, and both 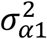 and 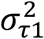 were set at 0.05. It is noted that the small-effects setting resembled more closely parameter estimates from real data analyses than the large-effects setting. Thus, results of the former setting would be more informative about how well our models and the likelihood test for model comparisons perform for analysis of real data.

Each parameter setting covered four scenarios—no G-C and R-C interactions (or the null), R-C interaction only, G-C interaction only, and both R-C and G-C interactions— where the true data generating models were the four models described above. Under each scenario, we simulated 100 replicates of phenotypic data (*n* = 7,513) of a main trait and a covariate, each based on 10,000 randomly chosen causal variants from the ARIC genotype data (see Supplementary Table 1 for an overview). For every replicate, we fitted the full and null models and compared the fit of the two models using the abovementioned likelihood ratio test. For every scenario, we computed the proportion of replicates, out of 100, for which the full model has a better fit than the null. This proportion takes on different interpretations depending on the simulation scenario. It is an estimate of type I error rate when the true model is the null, whereas it is an estimate of statistical power in scenarios where the true model is other than the null. It is important to note that all simulating models above assume normally distributed random effects (e.g., genetic and residual effects). In effect, for any given covariate value, the main trait follows a normal distribution. This normality assumption however, is likely violated for many traits of the ARIC and UKBB datasets, which are characterised by substantially larger kurtosis and skewness than would be expected from data simulated under normality (Supplementary Figure 7). Therefore, in addition to the large and small effects settings described above, we also simulated data with non-normal residuals drawn from Gamma distributions. We purposefully chose two sets of shape and scale parameters of Gamma distributions to represent large and small deviations from normality. For each of the non-normal settings, we had two scenarios: no G-C and R-C interaction (i.e., the null model is true) and G-C and R-C interactions (i.e., the full model is true), each with 100 replicates. We fitted the null and full models, which by definition all assume normality of random effects, to each replicate and subsequently assessed our model comparison method, in terms of type I error, power, and parameter estimates. In the event of an inflated type I error rate, we applied a rank-based inverse normal transformation to the simulated data and refitted the models. We then assessed the effectiveness of the transformation on reducing false positive findings and its potential consequences on statistical power of the model comparison method and model parameter estimates.

It is important to emphasize that our interaction models do not assume absolute normality of phenotype data, rather their conditional normality on the covariate. Thus, unless the true underlying model is the null, when phenotypic observations are collapsed across covariate values, the distribution of the collapsed data is not necessarily normal. In fact, in the presence of genuine G-C and/or R-C interactions, even when the model normality assumption is met, the simulated phenotype can have larger skewness and kurtosis than data simulated under the null model (see Supplementary Figure 7). Thus, deviations from normality of a given set of phenotype data could arise from genuine G-C or R-C interaction. If they are mistaken as signs of violation of the model normality assumption, what would be the consequences of applying a rank-based inverse normal transformation for type I error rate, statistical power and model estimates? To answer these questions, we also applied the transformation to phenotype data simulated under normality (i.e., large & small effect parameter settings) and assessed its impact on type I error rate, statistical power and model estimates.

### Validation

To validate significant results found in the ARIC dataset, we repeated analyses using the UKBB for variables where the two datasets overlap. Since the UKBB has a larger sample size, hence greater statistical power, we explicitly estimated the genetic variance of the covariate when fitting a MRNM, rather than fixing this parameter at zero as for the ARIC dataset. Subsequently, the degree of freedom used for the likelihood ratio test that compares the full model with the null model was seven as opposed to five. Same as for the ARIC dataset, we estimated heritability of each UKBB trait using two URNMS (i.e., null and interaction models) and the inclusion of a covariate was based on MRNM results.

## URLs

ARIC study: https://www2.cscc.unc.edu/aric/

UK Biobank: http://www.ukbiobank.ac.uk/

MTG2 for fitting reaction norm models:

https://sites.google.com/site/honglee0707/mtg2

### Data Availability

Simulated data used in this paper can be obtained from the authors upon request. Our access to the ARIC data was under the code phs000090, and access to the UK biobank data was approved by the UK biobank research ethics committee under the reference number 14575.

## Acknowledgments

This research is supported by the Australian National Health and Medical Research Council (1080157, 1087889) and the Australian Research Council (DP160102126, DP190100766, FT160100229). The authors would like to thank staff and participants of the ARIC study and the UK Biobank for their important contributions.

## References

1. WHO. Fact Sheet: Cardiovascular Diseases. Vol. 2018 (WHO, 2018).

2. Waken, R.J., de las Fuentes, L. & Rao, D.C. A review of the genetics of hypertension with a focus on gene-environment interactions. Current hypertension reports 19, 23 (2017).

3. Namboodiri, K.K. et al. The Collaborative Lipid Research Clinics Family Study: biological and cultural determinants of familial resemblance for plasma lipids and lipoproteins. Genetic epidemiology 2, 227–254 (1985).

4. Freeman, M.S., Mansfield, M.W., Barrett, J.H. & Grant, P.J. Genetic contribution to circulating levels of hemostatic factors in healthy families with effects of known genetic polymorphisms on heritability. Arteriosclerosis, thrombosis, and vascular biology 22, 506–510 (2002).

5. Vossen, C.Y. et al. A genetic basis for the interrelation of coagulation factors. Journal of Thrombosis and Haemostasis 5, 1930–1935 (2007).

6. Nowak-Göttl, U. et al. Genetics of hemostasis: differential effects of heritability and household components influencing lipid concentrations and clotting factor levels in 282 pediatric stroke families. Environmental Health Perspectives 116, 839 (2008).

7. Robinson, M.R. et al. Genotype–covariate interaction effects and the heritability of adult body mass index. Nature Genetics 49, 1174 (2017).

8. Moore, R. et al. A linear mixed-model approach to study multivariate gene–environment interactions. Nature Genetics 51, 180–186 (2019).

9. Young, A.I., Wauthier, F. & Donnelly, P. Multiple novel gene-by-environment interactions modify the effect of FTO variants on body mass index. Nature Communications 7, 12724 (2016).

10. Ni, G. et al. Genotype–covariate correlation and interaction disentangled by a whole-genome multivariate reaction norm model. Nature communications 10, 2239 (2019).

11. Maher, B. Personal genomes: The case of the missing heritability. Nature News 456, 18–21 (2008).

12. Manolio, T.A. et al. Finding the missing heritability of complex diseases. Nature 461, 747 (2009).

13. Hindy, G., Wiberg, F., Almgren, P., Melander, O. & Orho-Melander, M. Polygenic risk score for coronary heart disease modifies the elevated risk by cigarette smoking for disease incidence. Circulation: Genomic and Precision Medicine 11, e001856 (2018).

14. Khera, A.V. et al. Genetic Risk, Adherence to a Healthy Lifestyle, and Coronary Disease. N Engl J Med 375, 2349–2358 (2016).

15. Rutten-Jacobs, L.C. et al. Genetic risk, incident stroke, and the benefits of adhering to a healthy lifestyle: cohort study of 306 473 UK Biobank participants. BMJ 363, k4168 (2018).

16. Corella, D. et al. APOA2, dietary fat, and body mass index: replication of a gene-diet interaction in 3 independent populations. Archives of internal medicine 169, 1897–1906 (2009).

17. Latella, M.C. et al. Genetic variation of alcohol dehydrogenase type 1C (ADH1C), alcohol consumption, and metabolic cardiovascular risk factors: results from the IMMIDIET study. Atherosclerosis 207, 284–290 (2009).

18. Bernstein, M.S. et al. Physical activity may modulate effects of ApoE genotype on lipid profile. Arteriosclerosis, thrombosis, and vascular biology 22, 133–140 (2002).

19. Corella, D. et al. Environmental factors modulate the effect of the APOE genetic polymorphism on plasma lipid concentrations: ecogenetic studies in a Mediterranean Spanish population. Metabolism-Clinical and Experimental 50, 936–944 (2001).

20. Ruaño, G. et al. Apolipoprotein A1 genotype affects the change in high density lipoprotein cholesterol subfractions with exercise training. Atherosclerosis 185, 65–69 (2006).

21. Lynch, M. & Walsh, B. Genetics and analysis of quantitative traits, (Sinauer Sunderland, MA, 1998).

22. Clark, S.A. & van der Werf, J. Genomic best linear unbiased prediction (gBLUP) for the estimation of genomic breeding values. Methods Mol Biol 1019, 321–30 (2013).

23. Henderson, C.R. Best linear unbiased estimation and prediction under a selection model. Biometrics, 423–447 (1975).

24. Kaprio, J. Twins and the mystery of missing heritability: the contribution of gene-environment interactions. J Intern Med 272, 440–8 (2012).

25. Bentley, A.R. et al. Multi-ancestry genome-wide gene-smoking interaction study of 387,272 individuals identifies new loci associated with serum lipids. Nat Genet 51, 636–648 (2019).

26. Yang, J. et al. Common SNPs explain a large proportion of the heritability for human height. Nat Genet 42, 565 (2010).

27. Dahl, A., Cai, N., Flint, J. & Zaitlen, N. GxEMM: Extending linear mixed models to general gene-environment interactions. bioRxiv, 397638 (2018).

28. Arena, R. et al. Applying Precision Medicine to Healthy Living for the Prevention and Treatment of Cardiovascular Disease. Curr Probl Cardiol 43, 448–483 (2018).

29. Leopold, J.A. & Loscalzo, J. Emerging Role of Precision Medicine in Cardiovascular Disease. Circ Res 122, 1302–1315 (2018).

30. Investigators, A. The atherosclerosis risk in communit (aric) stui) y: design and objectwes. American journal of epidemiology 129, 687–702 (1989).

31. Papp, A., Hatzakis, H., Bracey, A. & Wu, K. ARICHemostasis Study-I. Development of a Blood Collection and Processing System Suitable for Multicenter Hemostatic Studies. Thrombosis and haemostasis 61, 015–019 (1989).

32. Seaman, C.D., George, K.M., Ragni, M. & Folsom, A.R. Association of von Willebrand factor deficiency with prevalent cardiovascular disease and asymptomatic carotid atherosclerosis: The Atherosclerosis Risk in Communities Study. Thrombosis research 144, 236 (2016).

33. Folsom, A.R., Nieto, F.J., Sorlie, P., Chambless, L.E. & Graham, D.Y. Helicobacter pylori seropositivity and coronary heart disease incidence. Circulation 98, 845–850 (1998).

34. O’Neal, W.T. et al. Association Between QT-Interval Components and Sudden Cardiac Death: The ARIC Study (Atherosclerosis Risk in Communities). Circ Arrhythm Electrophysiol 10(2017).

35. Decker, W.W. et al. Continuous 12-lead electrocardiographic monitoring in an emergency department chest pain unit: an assessment of potential clinical effect. Ann Emerg Med 41, 342–51 (2003).

36. Willett, W.C. et al. Reproducibility and validity of a semiquantitative food frequency questionnaire. American journal of epidemiology 122, 51–65 (1985).

37. Shekelle, R.B. et al. Diet, serum cholesterol, and death from coronary heart disease: the Western Electric Study. New England Journal of Medicine 304, 65–70 (1981).

38. Stamler, J. et al. Higher blood pressure in middle-aged American adults with less education— role of multiple dietary factors: the INTERMAP study. Journal of human hypertension 17, 655 (2003).

39. Baecke, J.A., Burema, J. & Frijters, J.E. A short questionnaire for the measurement of habitual physical activity in epidemiological studies. The American journal of clinical nutrition 36, 936–942 (1982).

40. Folsom, A.R. et al. Physical activity and incidence of coronary heart disease in middle-aged women and men. Medicine and science in sports and exercise 29, 901–909 (1997).

41. Bell, E.J., Lutsey, P.L., Windham, B.G. & Folsom, A.R. Physical activity and cardiovascular disease in African Americans in ARIC. Medicine and science in sports and exercise 45, 901 (2013).

42. Sudlow, C. et al. UK biobank: an open access resource for identifying the causes of a wide range of complex diseases of middle and old age. PLoS Med 12, e1001779 (2015).

43. Cassidy, S., Chau, J.Y., Catt, M., Bauman, A. & Trenell, M.I. Cross-sectional study of diet, physical activity, television viewing and sleep duration in 233 110 adults from the UK Biobank; the behavioural phenotype of cardiovascular disease and type 2 diabetes. BMJ open 6, e010038 (2016).

44. Lee, S.H. et al. Genetic relationship between five psychiatric disorders estimated from genome-wide SNPs. Nature genetics 45, 984 (2013).

45. Ripke, S. et al. Genome-wide association analysis identifies 13 new risk loci for schizophrenia. Nature genetics 45, 1150 (2013).

46. Lee, S.H. et al. Estimation of SNP heritability from dense genotype data. The American Journal of Human Genetics 93, 1151–1155 (2013).

47. Lee, S.H. & Van der Werf, J.H. MTG2: an efficient algorithm for multivariate linear mixed model analysis based on genomic information. Bioinformatics 32, 1420–1422 (2016).

