## Supplemental Materials for "A Whole-Genome Approach Discovers Novel Genetic and Non-Genetic Variance Components Modulated by Lifestyle for Cardiovascular Health"

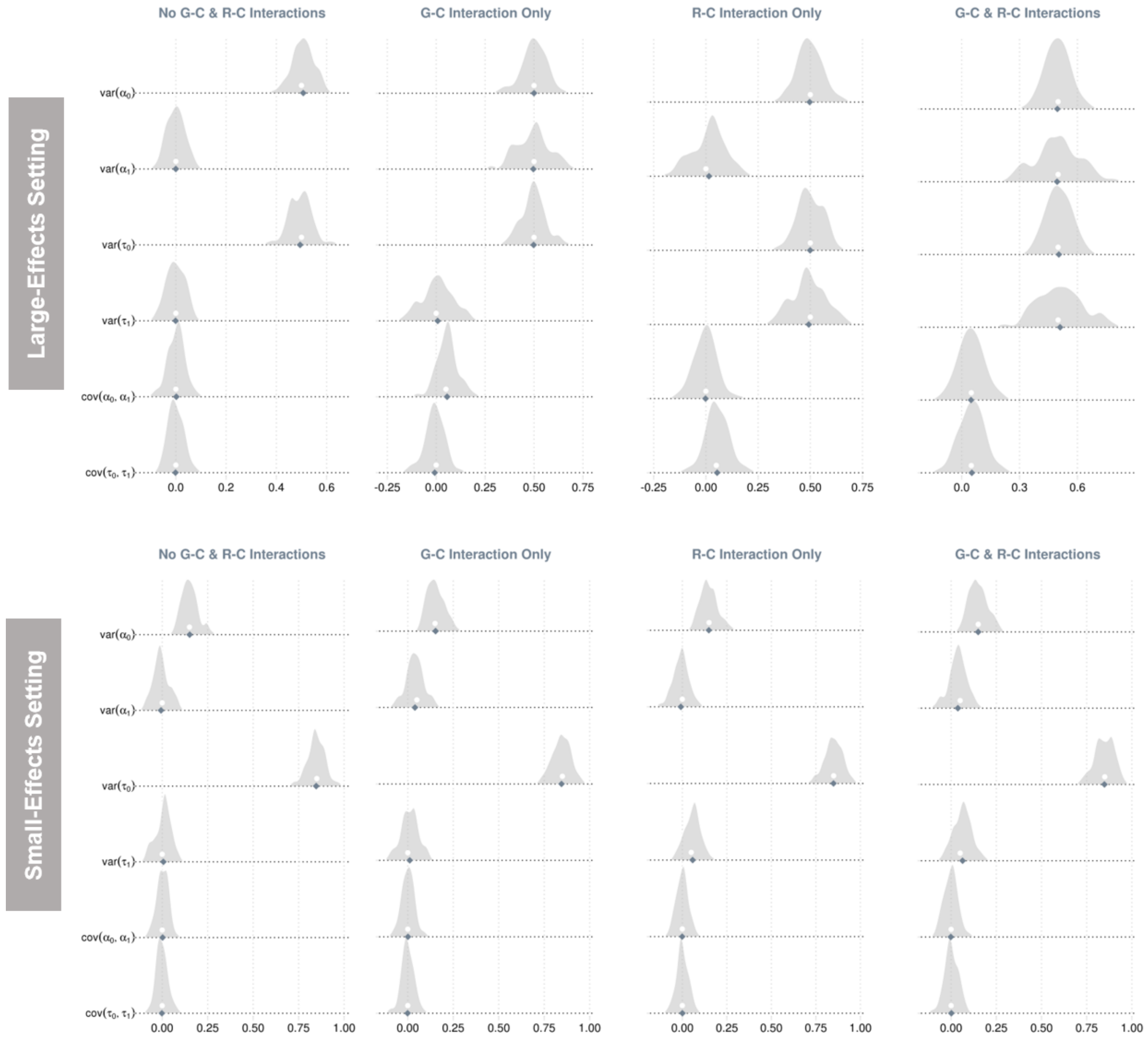


**Sup. Figure 1. Sampling distributions of parameter estimates from the full model.** Data for a main trait and a covariate were simulated using 4 multivariate reaction norm models under 2 parameter settings for each model, which together gave arise to 8 combinations of simulation scenarios. The four models were no G-C and R-C interactions (i.e., a null model), G-C interaction only (i.e., a G-C model), R-C interaction only (i.e., a R-C model), and both G-C and R-C interactions (i.e., a full model). The two parameter settings were large and small effect sizes in terms of heritability, G-C and R-C interactions. Each simulation was repeated 100 times, resulting in 100 replicates of simulated data under each scenario. Parameter estimates were obtained from fitting the full model. Shown distributions are for model parameters (i.e., variance & covariance terms) pertaining to the main trait only. True parameter values are shown in dots and means of sampling distributions in diamonds.

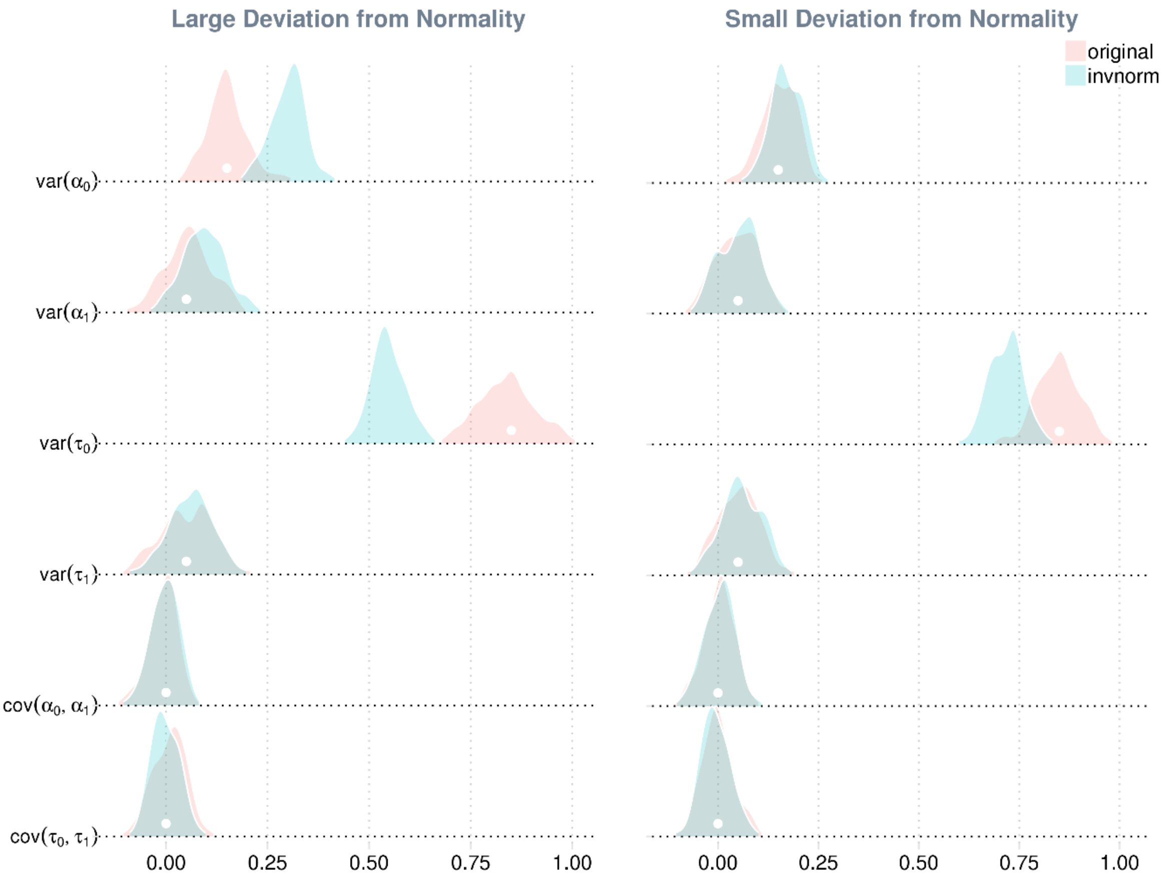


**Sup. Figure 2. Impact of rank-based inverse normal transformation on parameter estimates when the normality assumption is violated.** Data for a main trait and a covariate were simulated using a multivariate reaction norm model that included both G-C and R-C interactions (i.e., a full model) under two parameter settings, where residuals of the main trait were drawn from distributions that deviated from a normal distribution to different degrees (i.e., small vs. large deviation from normality). Each simulation was repeated 100 times, resulting in 100 replicates of simulated data for each setting. Parameter estimates were obtained from fitting a full model—that assumes normality of all random effects including residuals—to the simulated data before and after a rank-based inverse normal distribution ('original' vs. 'invnorm'). Shown distributions are for model parameters (i.e., variance & covariance terms) pertaining to the main trait only. True parameter values are shown in dots.

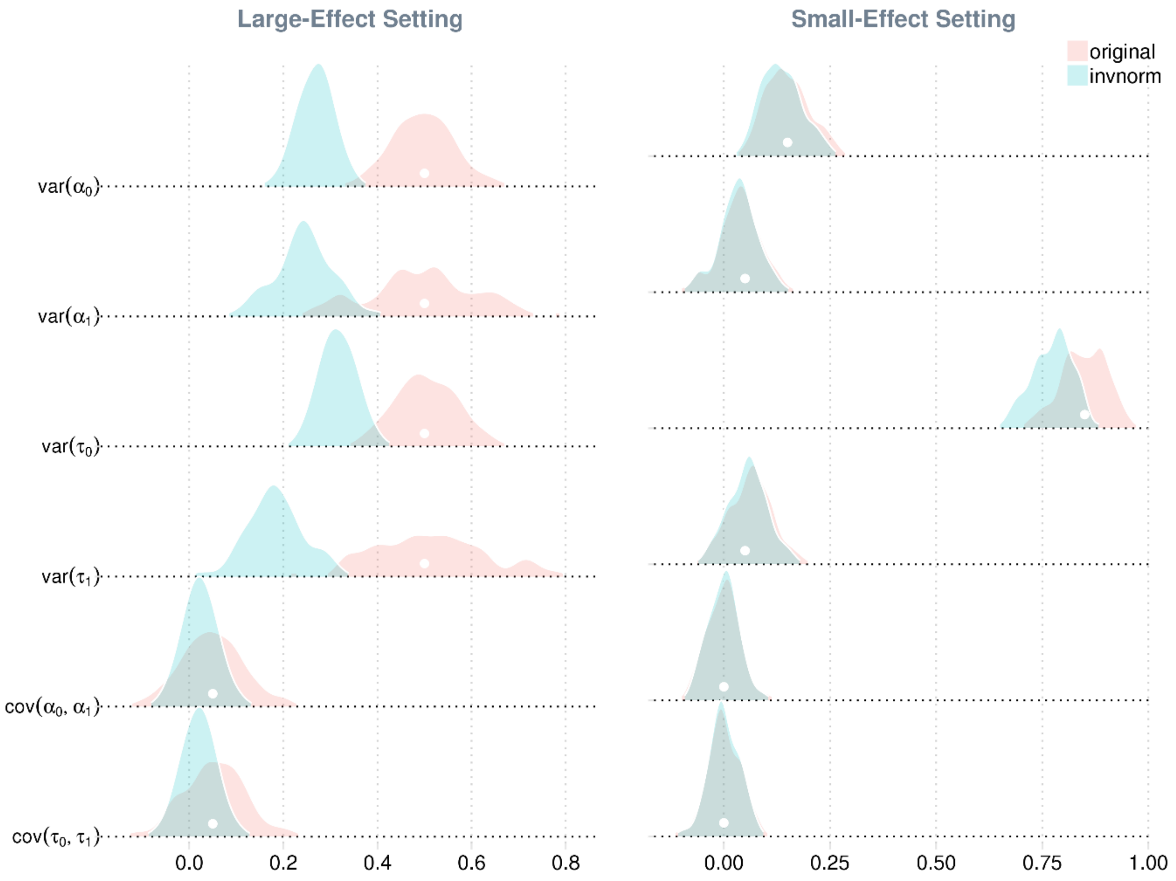


**Sup. Figure 3. Impact of rank-based inverse normal transformation on parameter estimates when normality assumption is met.** Data for a main trait and a covariate were simulated using a multivariate reaction norm model that included both G-C and R-C interactions (i.e., a full model) under two parameter settings that varied in effect sizes in terms of heritability, G-C and R-C interactions. In each setting, all random effects were drawn from normal distributions. Each simulation was repeated 100 times, resulting in 100 replicates of simulated data for each setting. Parameter estimates were obtained from fitting a full model—that assumes normality of random effects—to the simulated data before and after a rank-based inverse normal distribution ('original' vs. 'invnorm'). Shown distributions are for model parameters (i.e., variance & covariance terms) pertaining to the main trait only. True parameter values are shown in dots.

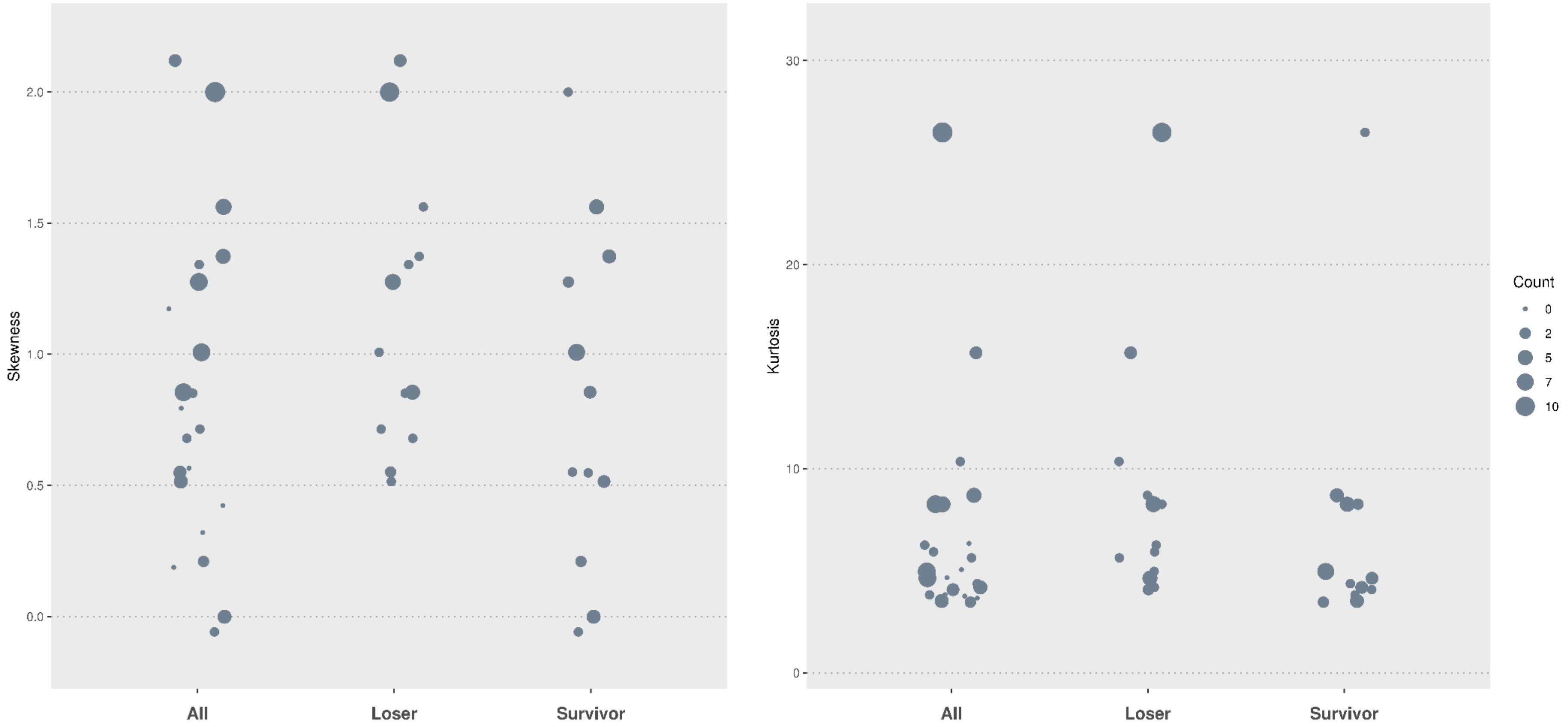


**Sup. Figure 4. Skewness (left panel) and kurtosis (right) of cardiovascular traits from the ARIC dataset by survivorship of rank-based inverse normal transformation.** Categories from left to right in each panel are all traits (labelled as "all"), trait that lost one or more signals after a rank-based inverse normal transformation ("loser"), and traits that still had one or more signals after the transformation ("survivor"). Point size is proportional to the count of signals. To reduce overlaps, points are jittered randomly in the horizontal direction. Note that a trait can be both a loser and a survivor.

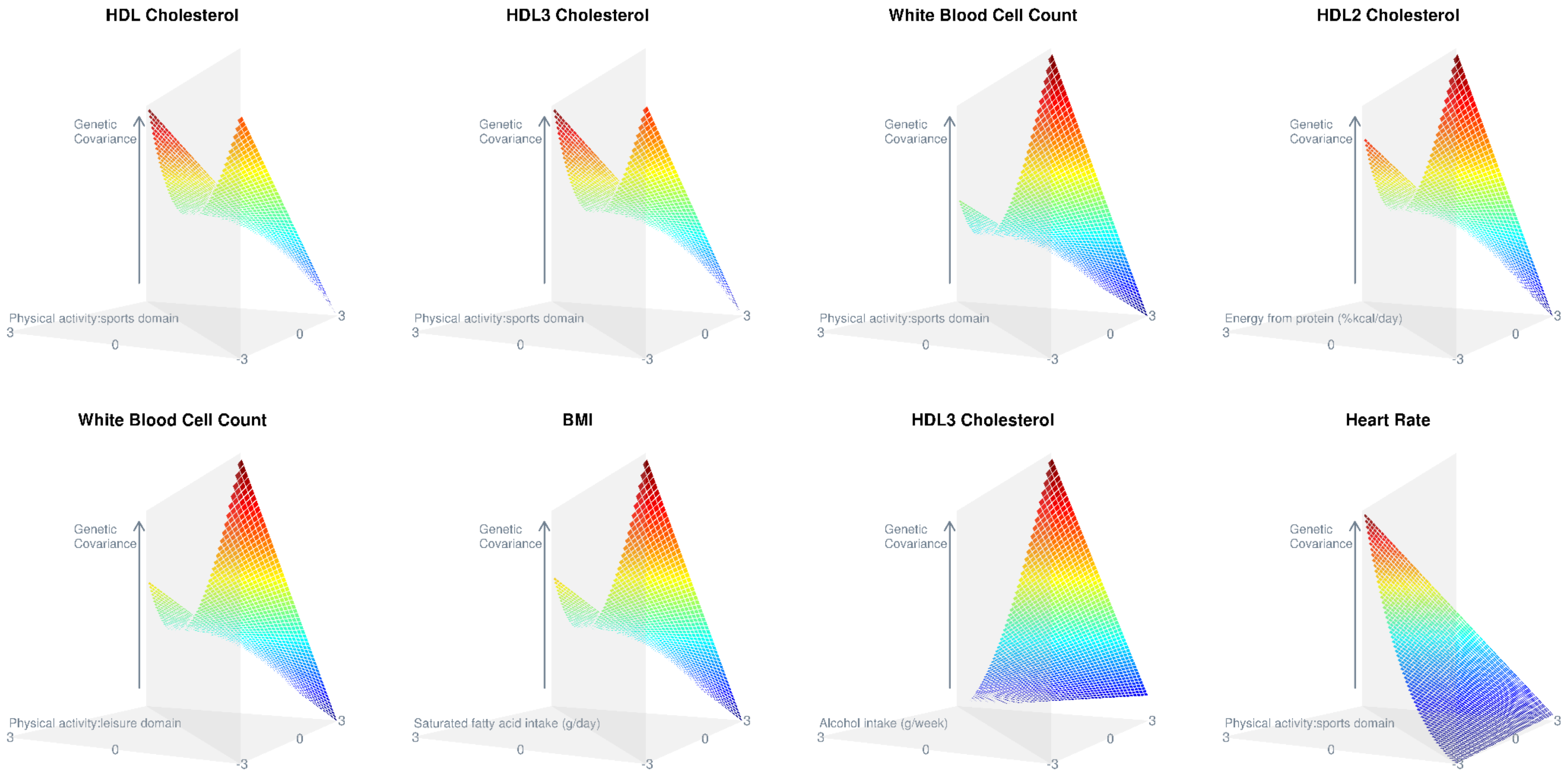


**Sup. Figure 5. Estimated genetic covariance with respect to lifestyle covariate.** Both x and y coordinates of each plot cover three standard deviations from the mean of a given lifestyle covariate. The horizontal plane thus represents pairwise combinations of lifestyle covariate values. The corresponding genetic covariance matrix of these combinations, estimated from the full model, is shown as the surface in each plot. The lower triangular part of each matrix, which is identical as the upper triangular part, is removed for simplicity. Diagonal entries of each matrix, shown as the intersection of the surface with the diagonal plane, are estimated genetic variances. Only traits with the first eight largest variance estimates of G-C interaction (see Figure 2 left) are shown. Arrows point to higher values.

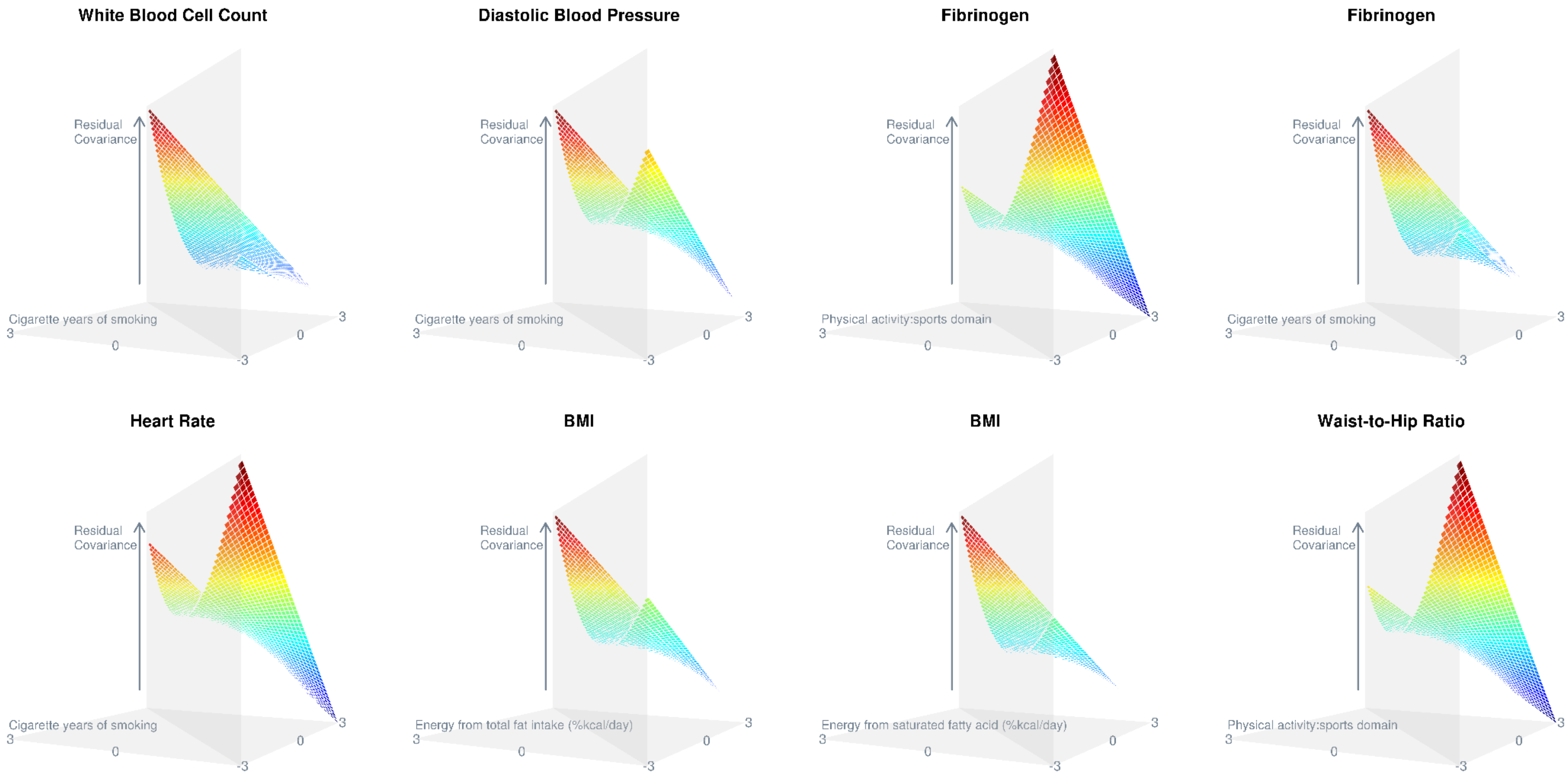


**Sup. Figure 6. Estimated residual covariance with respect to lifestyle covariate.** Both x and y coordinates of each plot cover three standard deviations from the mean of a given lifestyle covariate. The horizontal plane thus represents pairwise combinations of lifestyle covariate values. The corresponding residual covariance matrix of these combinations, estimated from the full model, is shown as the surface in each plot. The lower triangular part of each matrix, which is identical as the upper triangular part, is removed for simplicity. Diagonal entries of each matrix, shown as the intersection of the surface with the diagonal plane, are estimated residual variances. Only traits with the first eight largest variance estimates of R-C interaction (see Figure 2 right) are shown. Arrows point to higher values.

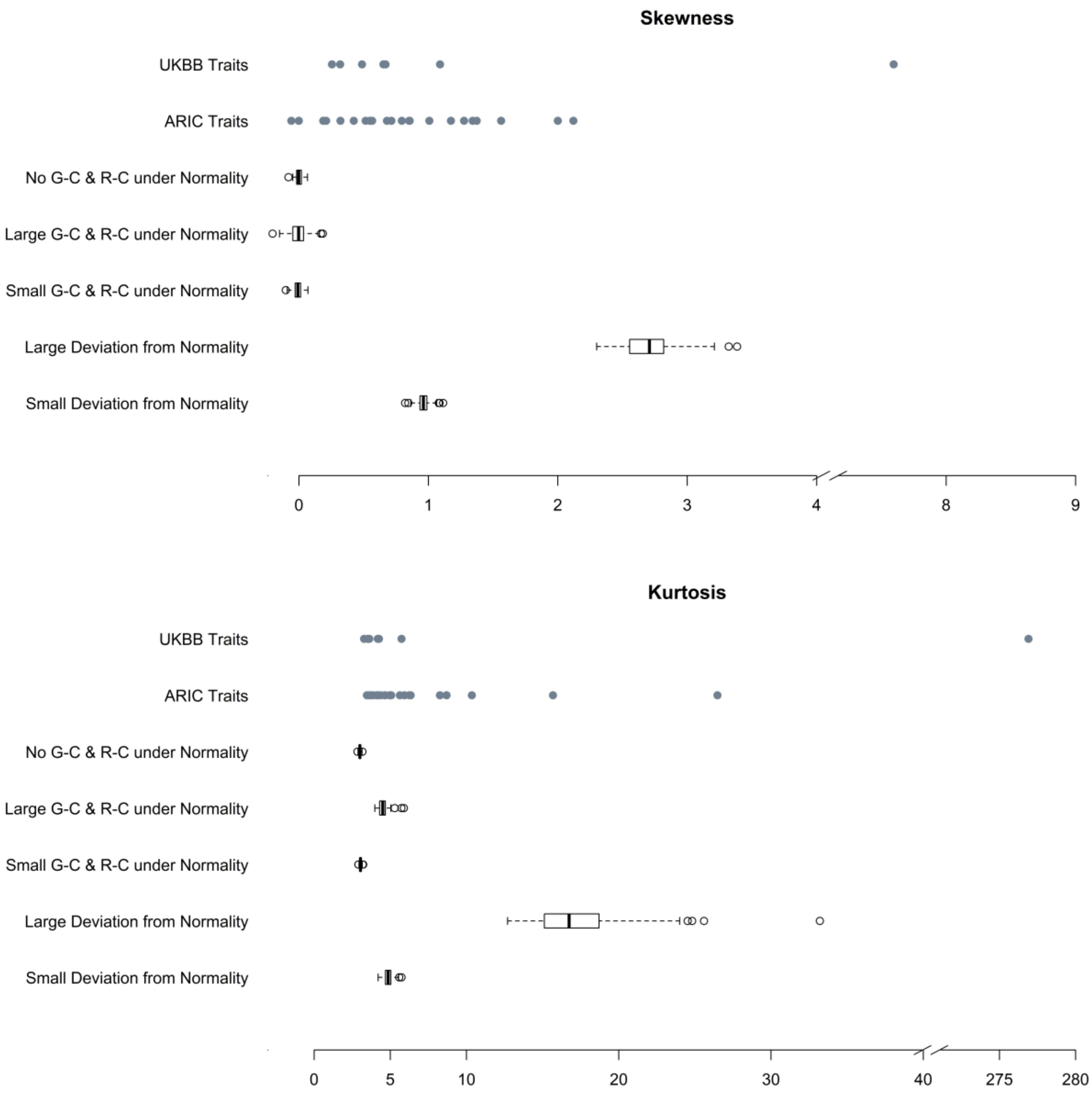


**Sup. Figure 7. Skewness (top) and kurtosis (bottom) of cardiovascular traits from the ARIC and UKBB datasets.** In each panel, categories along the y axis from the top to bottom are traits from the UK biobank, traits from the ARIC study, traits (100 replicates) simulated from the null model, from the full model with large and small effect sizes, and from the full model with non-normal residuals that result in large and small phenotypic deviations from normality. The last five categories are included as references to indicate expected skewness and kurtosis when the model assumption of normality is and is not met.

**Sup. Table 1. True parameter values of four simulation models under four settings.**

#### A. Small effects under Normality

| Parameter | No G-C & R-C | G-C & R-C | G-C only | R-C only |
|---|---|---|---|---|
| var($\alpha_0$) | 0.15 | 0.15 | 0.15 | 0.15 |
| var($\alpha_1$) | 0 | 0.05 | 0.05 | 0 |
| var($\tau_0$) | 0.85 | 0.85 | 0.85 | 0.85 |
| var($\tau_1$) | 0 | 0.05 | 0 | 0.05 |
| var($\beta$) | 0 | 0 | 0 | 0 |
| var($\varepsilon$) | 1 | 1 | 1 | 1 |
| cov($\alpha_0$, $\alpha_1$) | 0 | 0 | 0 | 0 |
| cov($\tau_0$, $\tau_1$) | 0 | 0 | 0 | 0 |
| cov($\alpha_0$, $\beta$) | 0 | 0 | 0 | 0 |
| cov($\alpha_1$, $\beta$) | 0 | 0 | 0 | 0 |
| cov($\tau_0$, $\varepsilon$) | 0 | 0 | 0 | 0 |
| cov($\tau_1$, $\varepsilon$) | 0 | 0 | 0 | 0 |

#### B. Large effects under Normality

| Parameter | No G-C & R-C | G-C & R-C | G-C only | R-C only |
|---|---|---|---|---|
| var($\alpha_0$) | 0.5 | 0.5 | 0.5 | 0.5 |
| var($\alpha_1$) | 0 | 0.5 | 0.5 | 0 |
| var($\tau_0$) | 0.5 | 0.5 | 0.5 | 0.5 |
| var($\tau_1$) | 0 | 0.5 | 0 | 0.5 |
| var($\beta$) | 0.5 | 0.5 | 0.5 | 0.5 |
| var($\varepsilon$) | 0.5 | 0.5 | 0.5 | 0.5 |
| cov($\alpha_0$, $\alpha_1$) | 0 | 0.05 | 0.05 | 0 |
| cov($\tau_0$, $\tau_1$) | 0 | 0.05 | 0 | 0.05 |
| cov($\alpha_0$, $\beta$) | 0 | 0 | 0 | 0 |
| cov($\alpha_1$, $\beta$) | 0 | 0 | 0 | 0 |
| cov($\tau_0$, $\varepsilon$) | 0 | 0 | 0 | 0 |
| cov($\tau_1$, $\varepsilon$) | 0 | 0 | 0 | 0 |

#### C. Small Deviation from Normality

| Parameter | No G-C & R-C | G-C & R-C | G-C only | R-C only |
|---|---|---|---|---|
| var($\alpha_0$) | 0.15 | 0.15 | 0.15 | 0.15 |
| var($\alpha_1$) | 0 | 0.05 | 0.05 | 0 |
| var($\tau_0$) | 0.85 | 0.85 | 0.85 | 0.85 |
| var($\tau_1$) | 0 | 0.05 | 0 | 0.05 |
| var($\beta$) | 0 | 0 | 0 | 0 |
| var($\varepsilon$) | 1 | 1 | 1 | 1 |
| cov($\alpha_0$, $\alpha_1$) | 0 | 0 | 0 | 0 |
| cov($\tau_0$, $\tau_1$) | 0 | 0 | 0 | 0 |
| cov($\alpha_0$, $\beta$) | 0 | 0 | 0 | 0 |
| cov($\alpha_1$, $\beta$) | 0 | 0 | 0 | 0 |
| cov($\tau_0$, $\varepsilon$) | 0 | 0 | 0 | 0 |
| cov($\tau_1$, $\varepsilon$) | 0 | 0 | 0 | 0 |
| $k_0$ | 2 | 2 | 2 | 2 |
| $\theta_0$ | 0.65 | 0.65 | 0.65 | 0.65 |
| $k_1$ | - | 2 | - | 2 |
| $\theta_1$ | - | 0.16 | - | 0.16 |

#### D. Large Deviation from Normality

| Parameter | No G-C & R-C | G-C & R-C | G-C only | R-C only |
|---|---|---|---|---|
| var($\alpha_0$) | 0.15 | 0.15 | 0.15 | 0.15 |
| var($\alpha_1$) | 0 | 0.05 | 0.05 | 0 |
| var($\tau_0$) | 0.85 | 0.85 | 0.85 | 0.85 |
| var($\tau_1$) | 0 | 0.05 | 0 | 0.05 |
| var($\beta$) | 0 | 0 | 0 | 0 |
| var($\varepsilon$) | 1 | 1 | 1 | 1 |
| cov($\alpha_0$, $\alpha_1$) | 0 | 0 | 0 | 0 |
| cov($\tau_0$, $\tau_1$) | 0 | 0 | 0 | 0 |
| cov($\alpha_0$, $\beta$) | 0 | 0 | 0 | 0 |
| cov($\alpha_1$, $\beta$) | 0 | 0 | 0 | 0 |
| cov($\tau_0$, $\varepsilon$) | 0 | 0 | 0 | 0 |
| cov($\tau_1$, $\varepsilon$) | 0 | 0 | 0 | 0 |
| $k_0$ | 0.25 | 0.25 | 0.25 | 0.25 |
| $\theta_0$ | 1.84 | 1.84 | 1.84 | 1.84 |
| $k_1$ | - | 0.25 | - | 0.25 |
| $\theta_1$ | - | 0.45 | - | 0.45 |

The top two settings are under the assumption that all random effects of the multivariate reaction normal models (MRNMs) for simulation are drawn from normal distributions. This assumption is relaxed for the bottom two settings, where residual effects, $\tau_0$ and $\tau_1$, are drawn from Gamma($k_0$, $\theta_0$) and Gamma($k_1$, $\theta_1$) with mean centred at zero, respectively. Each setting comprises four simulation models, which from the left to right are the null, full, G-C and R-C models.

**Sup. Table 2. Proportion of simulated replicates for which the full model had a better fit than the null under different simulation scenarios.**

| Simulation model | Simulation under Normality | | | | Simulation under Non-Normality | | | | Interpretation |
|---|---|---|---|---|---|---|---|---|---|
| | large effects | | small effects | | large deviation | | small deviation | | |
| | *no RINT** | *RINT* | *no RINT* | *RINT* | *no RINT* | *RINT* | *no RINT* | *RINT* | |
| No G-C & R-C | 0.04 | 0.04 | 0.04 | 0.05 | 0.65 | 0.05 | 0.2 | 0.07 | type I error |
| G-C only | 1 | 1 | 0.93 | 0.94 | 0.81 | 1 | 0.84 | 0.99 | power |
| R-C only | 1 | 1 | 0.88 | 0.87 | 0.84 | 1 | 0.87 | 0.99 | power |
| G-C & R-C | 1 | 1 | 1 | 1 | 1 | 1 | 1 | 1 | power |

*RINT = Rank-based Inverse Normal Transformation

Data of a main trait and a covariate were simulated using four models ($1^{st}$ column) under normality with large and small effects (in terms of heritability, G-C and R-C interactions) and under non-normality that resulted in large and small phenotypic deviations of the main trait from normality. Each simulation was repeated 100 times, resulting in 100 replicates of simulated data under each setting. For each replicate, the full model, which allows G-C and R-C interactions, and the null model, which assumes no G-C and R-C interactions, were fitted then compared using a likelihood ratio test. The model comparison was repeated after a rank-based inverse normal transformation was applied to the simulated data.

**Sup. Table 3. Variance and covariance estimates from the full model for the 34 signals emerged from the ARIC dataset.**

| Main Trait | Lifestlye Covariate | var($\alpha_0$) | var($\alpha_1$) | cov($\alpha_0, \alpha_1$) | var($\tau_0$) | var($\tau_1$) | cov($\tau_0, \tau_1$) |
|---|---|---|---|---|---|---|---|
| Fibrinogen | Cigarette years of smoking | 4.0e+02(1.5e+02) | -3.6e+02(1.5e+02) | -7.9e+01(1.1e+02) | 3.1e+03(1.6e+02) | 2.9e+02(1.6e+02) | 3.6e+02(1.2e+02) |
| Fibrinogen | Physical activity:sports domain | 4.2e+02(1.5e+02) | -2.5e+02(1.3e+02) | 1.5e+01(1.0e+02) | 2.9e+03(1.6e+02) | 3.6e+02(1.4e+02) | -2.7e+02(1.1e+02) |
| Factor VII | Physical activity:sports domain | 1.0e+02(3.6e+01) | 2.6e+00(3.1e+01) | -1.5e+01(2.4e+01) | 6.5e+02(3.8e+01) | 7.2e+00(3.2e+01) | -4.9e+01(2.5e+01) |
| BMI | Alcohol intake (g/week) | 2.9e+00(9.0e-01) | 1.8e-01(6.9e-01) | -1.1e+00(5.7e-01) | 1.7e+01(9.3e-01) | 5.7e-01(6.9e-01) | -1.3e+00(6.0e-01) |
| BMI | Physical activity:leisure domain | 3.2e+00(9.2e-01) | -6.7e-02(8.3e-01) | 3.1e-01(6.3e-01) | 1.7e+01(9.6e-01) | 5.9e-01(8.7e-01) | -1.3e+00(6.4e-01) |
| BMI | Physical activity:sports domain | 3.1e+00(9.1e-01) | 1.1e-01(6.6e-01) | 2.3e-02(5.8e-01) | 1.7e+01(9.6e-01) | 4.2e-01(7.1e-01) | -2.5e+00(6.1e-01) |
| BMI | Keys score | 3.2e+00(9.2e-01) | -4.0e-01(8.0e-01) | -3.3e-01(6.2e-01) | 1.7e+01(9.6e-01) | 5.5e-01(8.3e-01) | 1.5e+00(6.3e-01) |
| BMI | Saturated fat intake (g/day) | 3.1e+00(9.2e-01) | 4.3e-01(8.7e-01) | -2.5e-01(6.5e-01) | 1.7e+01(9.6e-01) | -5.8e-01(9.1e-01) | 1.2e+00(6.6e-01) |
| BMI | Energy from saturated fat (%kcal/day) | 3.1e+00(9.2e-01) | -1.2e+00(8.1e-01) | -5.2e-01(6.2e-01) | 1.7e+01(9.6e-01) | 1.5e+00(8.5e-01) | 1.6e+00(6.4e-01) |
| BMI | Energy from total fat intake (%kcal/day) | 3.1e+00(9.2e-01) | -1.2e+00(8.2e-01) | -3.4e-01(6.3e-01) | 1.7e+01(9.6e-01) | 1.5e+00(8.8e-01) | 1.3e+00(6.4e-01) |
| Waist-to-Hip Ratio | Cigarette years of smoking | 3.0e-04(2.0e-04) | 0.0e+00(2.0e-04) | -1.0e-04(1.0e-04) | 3.7e-03(2.0e-04) | 0.0e+00(2.0e-04) | -1.0e-04(1.0e-04) |
| Waist-to-Hip Ratio | Physical activity:sports domain | 3.0e-04(2.0e-04) | -2.0e-04(1.0e-04) | 0.0e+00(1.0e-04) | 3.8e-03(2.0e-04) | 3.0e-04(2.0e-04) | -2.0e-04(1.0e-04) |
| Waist-to-Hip Ratio | Total energy intake (kcal/day) | 3.0e-04(2.0e-04) | -1.0e-04(1.0e-04) | 1.0e-04(1.0e-04) | 3.9e-03(2.0e-04) | 0.0e+00(2.0e-04) | -3.0e-04(1.0e-04) |
| Waist-to-Hip Ratio | Energy from protein intake (%kcal/day) | 3.0e-04(2.0e-04) | -1.0e-04(2.0e-04) | -1.0e-04(1.0e-04) | 3.8e-03(2.0e-04) | 0.0e+00(2.0e-04) | 2.0e-04(1.0e-04) |
| Pulse Pressure | Cigarette years of smoking | 8.0e+00(5.1e+00) | -1.1e+00(5.5e+00) | -2.7e+00(3.8e+00) | 1.0e+02(5.5e+00) | 6.3e+00(5.8e+00) | 4.6e+00(4.1e+00) |
| Diastolic Blood Pressure | Cigarette years of smoking | 7.4e+00(3.2e+00) | -7.3e+00(3.0e+00) | -2.5e+00(2.2e+00) | 6.4e+01(3.4e+00) | 8.1e+00(3.2e+00) | 4.4e+00(2.4e+00) |
| Heart Rate | Cigarette years of smoking | 1.3e+01(4.1e+00) | -2.1e+00(4.3e+00) | 1.9e+00(3.0e+00) | 7.1e+01(4.3e+00) | 6.8e+00(4.6e+00) | -1.6e+00(3.1e+00) |
| Heart Rate | Physical activity:leisure domain | 1.3e+01(4.1e+00) | -4.4e+00(3.4e+00) | 4.9e+00(2.7e+00) | 7.4e+01(4.3e+00) | 5.9e+00(3.7e+00) | -8.8e+00(2.8e+00) |
| Heart Rate | Physical activity:sports domain | 1.2e+01(4.0e+00) | 1.1e+00(3.2e+00) | 3.4e+00(2.6e+00) | 7.7e+01(4.3e+00) | -4.6e-01(3.4e+00) | -6.7e+00(2.7e+00) |
| HDL2 Cholesterol | Cigarette years of smoking | -1.0e-04(1.8e-03) | -3.0e-04(1.3e-03) | 5.0e-04(1.1e-03) | 4.0e-02(1.9e-03) | 4.0e-04(1.4e-03) | -3.1e-03(1.1e-03) |
| HDL2 Cholesterol | Physical activity:leisure domain | -6.0e-04(1.8e-03) | -1.2e-03(1.7e-03) | 2.0e-04(1.2e-03) | 4.0e-02(1.9e-03) | 1.7e-03(1.8e-03) | 3.0e-03(1.3e-03) |
| HDL2 Cholesterol | Carbohydrate intake (g/day) | -5.0e-04(1.8e-03) | -1.1e-03(1.4e-03) | -4.0e-04(1.1e-03) | 4.0e-02(1.9e-03) | 1.2e-03(1.4e-03) | -1.9e-03(1.1e-03) |
| HDL2 Cholesterol | Total energy intake (kcal/day) | -4.0e-04(1.8e-03) | -1.5e-03(1.3e-03) | -6.0e-04(1.1e-03) | 4.1e-02(1.9e-03) | 6.0e-04(1.4e-03) | -9.0e-04(1.2e-03) |
| HDL2 Cholesterol | Energy from protein intake (%kcal/day) | -4.0e-04(1.8e-03) | 1.6e-03(1.6e-03) | -4.0e-04(1.2e-03) | 4.1e-02(1.9e-03) | -2.2e-03(1.6e-03) | 2.4e-03(1.3e-03) |
| HDL3 Cholesterol | Alcohol intake (g/week) | 5.9e-03(2.7e-03) | 1.0e-03(3.3e-03) | -3.9e-03(2.1e-03) | 5.5e-02(2.9e-03) | -7.0e-04(3.3e-03) | 5.9e-03(2.3e-03) |
| HDL3 Cholesterol | Physical activity:sports domain | 6.2e-03(2.7e-03) | 5.0e-03(2.8e-03) | 1.4e-03(2.0e-03) | 5.1e-02(2.9e-03) | -1.8e-03(2.9e-03) | -2.4e-03(2.0e-03) |
| HDL Cholesterol | Physical activity:sports domain | 1.4e-02(6.1e-03) | 1.4e-02(6.3e-03) | 4.8e-03(4.3e-03) | 1.2e-01(6.6e-03) | -7.8e-03(6.4e-03) | -7.0e-03(4.4e-03) |
| HDL Cholesterol | Monounsaturated fatty acid intake (g/day) | 1.3e-02(6.2e-03) | 9.0e-04(5.2e-03) | -2.0e-03(4.0e-03) | 1.2e-01(6.5e-03) | -9.0e-04(5.2e-03) | -4.4e-03(4.2e-03) |
| HDL Cholesterol | Energy from protein intake (%kcal/day) | 1.3e-02(6.2e-03) | -4.7e-03(5.9e-03) | 1.0e-03(4.4e-03) | 1.2e-01(6.6e-03) | 5.8e-03(6.1e-03) | 4.3e-03(4.5e-03) |
| Apolipoprotein AI | Polyunsaturated fatty acid intake (g/day) | 7.3e+03(3.4e+03) | -3.4e+03(2.9e+03) | 3.7e+03(2.1e+03) | 6.8e+04(3.5e+03) | 3.8e+03(2.9e+03) | -7.4e+03(2.3e+03) |
| White Blood Cell Count | Cigarette years of smoking | 4.1e-01(1.2e-01) | -6.6e-01(4.6e-02) | -3.0e-01(8.0e-02) | 3.0e+00(1.4e-01) | 5.8e-01(7.4e-02) | 8.6e-01(9.2e-02) |
| White Blood Cell Count | Physical activity:leisure domain | 5.7e-01(1.3e-01) | 1.1e-01(1.3e-01) | -7.1e-02(9.4e-02) | 2.3e+00(1.4e-01) | 1.2e-02(1.3e-01) | -1.7e-01(9.5e-02) |
| White Blood Cell Count | Physical activity:sports domain | 5.5e-01(1.3e-01) | 1.4e-01(1.3e-01) | -1.3e-01(9.5e-02) | 2.2e+00(1.4e-01) | 9.1e-02(1.4e-01) | -9.1e-02(9.7e-02) |
| White Blood Cell Count | Keys score | 5.5e-01(1.3e-01) | 3.9e-03(1.2e-01) | 1.5e-01(9.3e-02) | 2.4e+00(1.4e-01) | -1.0e-02(1.3e-01) | 3.0e-03(9.3e-02) |

All estimates are derived from analyses of data without a rank-based inverse normal transformation. Standard errors are in brackets. Other model parameters are omitted for simplicity. Signals replicated in the UK biobank are shaded. Note the UK biobank only has data available to validate 17 signals emerged from ARIC.

**Sup. Table 4. Variance and covariance estimates from the full model for the UK Biobank dataset.**

| Main Trait | Lifestlye Covariate | var($\alpha_0$) | var($\alpha_1$) | cov($\alpha_0,\alpha_1$) | var($\tau_0$) | var($\tau_1$) | cov($\tau_0,\tau_1$) |
|---|---|---|---|---|---|---|---|
| BMI | Alcohol intake (glass & pint/week) | 3.9e+00(2.0e-01) | 4.4e-01(2.1e-01) | -1.4e-01(1.4e-01) | 1.3e+01(2.1e-01) | 3.2e-01(2.3e-01) | -4.4e-01(1.6e-01) |
| BMI | MET minutes/week for walking | 4.0e+00(2.0e-01) | 5.7e-02(1.7e-01) | -3.6e-01(1.3e-01) | 1.2e+01(2.3e-01) | 1.1e+00(1.9e-01) | -1.2e+00(1.5e-01) |
| BMI | MET minutes/week for moderate activity | 3.9e+00(2.0e-01) | -1.8e-01(1.4e-01) | -1.9e-01(1.2e-01) | 1.2e+01(2.2e-01) | 1.4e+00(1.8e-01) | -1.7e+00(1.5e-01) |
| BMI | MET minutes/week for vigorous activity | 4.0e+00(2.1e-01) | 1.6e-01(5.7e-01) | -1.6e-01(5.7e-01) | 1.3e+01(2.2e-01) | 5.8e-01(5.8e-01) | -1.8e+00(5.7e-01) |
| BMI | Summed MET minutes/week for all activity | 3.9e+00(3.7e-01) | -1.1e-01(1.9e-01) | -2.7e-01(2.4e-01) | 1.3e+01(2.1e-01) | 1.2e+00(1.9e-01) | -1.7e+00(2.2e-01) |
| BMI | estimated saturated fat intake | 3.9e+00(3.7e-01) | 4.7e-01(3.6e-01) | -9.6e-02(2.6e-01) | 1.3e+01(3.8e-01) | -2.1e-01(3.7e-01) | 1.9e-01(2.6e-01) |
| Diastolic Blood Pressure[1] | Pack years adult smoking as proportion of life span exposed to smoking | 1.6e+01(1.2e+00) | 2.2e+00(9.5e-01) | -1.8e+00(7.8e-01) | 9.1e+01(1.3e+00) | -2.7e+00(9.0e-01) | 3.2e+00(8.5e-01) |
| Pulse Pressure[1] | Pack years adult smoking as proportion of life span exposed to smoking | 2.5e+01(2.0e+00) | -9.5e-01(1.8e+00) | 2.1e+00(1.3e+00) | 1.5e+02(2.2e+00) | 2.4e-02(1.7e+00) | 2.0e+00(1.5e+00) |
| Heart Rate | Pack years adult smoking as proportion of life span exposed to smoking | 2.0e+01(1.5e+00) | 6.0e-01(1.5e+00) | -6.7e-01(1.1e+00) | 1.1e+02(1.8e+00) | 2.1e+00(1.7e+00) | 3.3e+00(1.2e+00) |
| Heart Rate | MET minutes/week for walking | 2.0e+01(1.5e+00) | 7.5e-01(1.3e+00) | 2.0e-01(9.7e-01) | 1.1e+02(1.8e+00) | 1.5e+00(1.5e+00) | -3.8e+00(1.2e+00) |
| Heart Rate | Summed MET minutes/week for all activity | 2.0e+01(1.5e+00) | -6.8e-01(1.3e+00) | -6.6e-02(1.0e+00) | 1.1e+02(1.7e+00) | 4.7e+00(1.5e+00) | -4.5e+00(1.2e+00) |
| Waist-to-Hip Ratio[2] | Pack years adult smoking as proportion of life span exposed to smoking | 1.5e-01(1.1e-02) | -3.6e-03(1.0e-02) | -4.7e-03(7.7e-03) | 8.5e-01(1.3e-02) | 4.1e-02(1.2e-02) | 3.4e-03(9.1e-03) |
| Waist-to-Hip Ratio | MET minutes/week for moderate activity | 1.5e-01(1.1e-02) | 5.6e-03(1.0e-02) | -1.2e-02(7.3e-03) | 8.1e-01(1.3e-02) | 2.9e-02(1.2e-02) | -2.3e-02(9.6e-03) |
| Waist-to-Hip Ratio | MET minutes/week for vigorous activity | 1.5e-01(1.1e-02) | 3.6e-03(1.0e-02) | -1.3e-02(7.5e-03) | 8.3e-01(1.2e-02) | 1.8e-02(1.1e-02) | -2.5e-02(9.4e-03) |
| Waist-to-Hip Ratio | Summed MET minutes/week for all activity | 1.5e-01(1.1e-02) | 9.5e-03(1.0e-02) | -1.1e-02(7.5e-03) | 8.2e-01(1.3e-02) | 1.9e-02(1.1e-02) | -1.9e-02(8.9e-03) |
| Waist-to-Hip Ratio | estimated total energy intake | 1.8e-01(2.2e-02) | -2.0e-02(2.0e-02) | 5.0e-03(1.5e-02) | 8.1e-01(2.3e-02) | 3.3e-02(2.1e-02) | -1.6e-02(1.5e-02) |
| White Blood Cell Count | Pack years adult smoking as proportion of life span exposed to smoking | 5.2e-01(4.2e-02) | -2.7e-02(4.2e-02) | -2.6e-02(3.0e-02) | 3.0e+00(5.2e-02) | 4.1e-01(5.4e-02) | -1.4e-01(3.6e-02) |
| HDL Cholesterol | MET minutes/week for moderate activity | 2.8e-02(1.4e-03) | 1.8e-03(1.3e-03) | 5.0e-04(1.0e-03) | 8.2e-02(1.6e-03) | -2.4e-03(1.4e-03) | 2.4e-03(1.2e-03) |
| HDL Cholesterol | MET minutes/week for vigorous activity | 2.8e-02(1.4e-03) | 2.0e-04(1.2e-03) | 1.0e-03(1.0e-03) | 8.1e-02(1.5e-03) | 7.0e-04(1.3e-03) | -1.2e-03(1.1e-03) |
| HDL Cholesterol | Summed MET minutes/week for all activity | 2.8e-02(1.4e-03) | 6.0e-04(1.2e-03) | 5.0e-04(1.0e-03) | 8.2e-02(1.5e-03) | -5.0e-04(1.3e-03) | 1.6e-03(1.1e-03) |

1.Estimation for the multivariate analysis did not converge. Shown are estimates from an univariate analysis, where the full model had a better fit than the null.

2.Estimates are based on standardized data for this trait, due to small phenotypic variance, which resulted in rather small estimates of variance components in absolute terms.

All estimates are derived from analyses of data without a rank-based inverse normal transformation. Standard errors are in brackets. Other model parameters are omitted for simplicity.

**Sup. Table 5. P-values for comparisons between the full model and nested models for the ARIC dataset.**

| Main Trait | Lifestlye Covariate | Null vs. Full | GC vs. Full | RC vs. Full |
|---|---|---|---|---|
| Fibrinogen | Cigarette years of smoking | 4.22E-10 | 1.83E-03 | 3.53E-01 |
| Fibrinogen | Physical activity:sports domain | 3.27E-06 | 2.57E-03 | 2.52E-01 |
| Factor VII | Physical activity:sports domain | 2.99E-05 | 7.19E-01 | 8.62E-01 |
| BMI | Alcohol intake (g/week) | 2.60E-12 | 5.80E-02 | 1.83E-01 |
| BMI | Physical activity:leisure domain | 9.04E-05 | 3.29E-01 | 8.69E-01 |
| BMI | Physical activity:sports domain | 9.23E-39 | 3.30E-08 | 9.60E-01 |
| BMI | Keys score | 1.28E-10 | 1.45E-01 | 9.73E-01 |
| BMI | Saturated fat intake (g/day) | 5.44E-05 | 4.66E-01 | 9.75E-01 |
| BMI | Energy from saturated fat (%kcal/day) | 6.24E-10 | 5.85E-02 | 3.72E-01 |
| BMI | Energy from total fat intake (%kcal/day) | 4.09E-08 | 9.83E-02 | 3.48E-01 |
| Waist-to-Hip Ratio | Cigarette years of smoking | 1.46E-09 | 1.34E-03 | 6.44E-01 |
| Waist-to-Hip Ratio | Physical activity:sports domain | 5.55E-10 | 2.35E-05 | 2.26E-01 |
| Waist-to-Hip Ratio | Total energy intake (kcal/day) | 2.00E-05 | 2.24E-01 | 4.35E-01 |
| Waist-to-Hip Ratio | Energy from protein (%kcal/day) | 7.11E-07 | 6.29E-04 | 7.63E-01 |
| Pulse Pressure | Cigarette years of smoking | 3.08E-06 | 2.50E-03 | 8.35E-01 |
| Diastolic Blood Pressure | Cigarette years of smoking | 5.43E-05 | 3.52E-04 | 5.93E-02 |
| Heart Rate | Cigarette years of smoking | 1.25E-06 | 1.37E-02 | 6.59E-01 |
| Heart Rate | Physical activity:leisure domain | 1.01E-04 | 8.47E-03 | 2.05E-01 |
| Heart Rate | Physical activity:sports domain | 3.74E-06 | 6.20E-06 | 4.18E-01 |
| HDL2 Cholesterol | Cigarette years of smoking | 1.70E-09 | 4.02E-02 | 6.94E-01 |
| HDL2 Cholesterol | Physical activity:leisure domain | 2.73E-12 | 1.32E-01 | 5.20E-01 |
| HDL2 Cholesterol | Carbohydrate intake (g/day) | 1.68E-07 | 3.42E-01 | 5.77E-01 |
| HDL2 Cholesterol | Total energy intake (kcal/day) | 2.40E-08 | 2.45E-01 | 4.50E-01 |
| HDL2 Cholesterol | Energy from protein (%kcal/day) | 3.69E-08 | 7.33E-02 | 6.54E-01 |
| HDL3 Cholesterol | Alcohol intake (g/week) | 3.31E-05 | 2.45E-02 | 1.88E-01 |
| HDL3 Cholesterol | Physical activity:sports domain | 9.68E-06 | 1.16E-03 | 1.52E-01 |
| HDL Cholesterol | Physical activity:sports domain | 8.85E-06 | 9.91E-05 | 4.71E-02 |
| HDL Cholesterol | Monounsaturated fatty acid intake (g/day) | 3.17E-05 | 4.31E-01 | 9.37E-01 |
| HDL Cholesterol | Energy from protein (%kcal/day) | 4.71E-06 | 7.86E-02 | 6.00E-01 |
| Apolipoprotein AI | Polyunsaturated fatty acid intake (g/day) | 2.94E-05 | 2.14E-02 | 2.13E-01 |
| White Blood Cell Count | Cigarette years of smoking | 2.00E-65 | 2.53E-21 | 1.25E-08 |
| White Blood Cell Count | Physical activity:leisure domain | 7.62E-07 | 5.19E-01 | 4.36E-01 |
| White Blood Cell Count | Physical activity:sports domain | 9.62E-05 | 1.72E-01 | 3.82E-01 |
| White Blood Cell Count | Keys score | 7.56E-05 | 8.37E-01 | 2.32E-01 |

Only shown for analyses where the full versus null model comparison was significant. Nested models include the null, G-C only and R-C only models. All analyses are based on data after a rank-based inverse normal transformation.

**Sup. Table 6. P-values for comparisons between the full model and nested models for the UK Biobank.**

| Main Trait | Lifestlye Covariate | Null vs. Full | GC vs. Full | RC vs. Full |
|---|---|---|---|---|
| BMI | Alcohol intake (glass & pint/week) | 1.19E-14 | 0.00E+00 | 0.00E+00 |
| BMI | MET minutes/week for walking | 1.49E-16 | 4.21E-04 | 6.82E-02 |
| BMI | MET minutes/week for moderate activity | 2.05E-29 | 6.34E-13 | 8.26E-01 |
| BMI | MET minutes/week for vigorous activity | 1.01E-46 | 8.91E-20 | 4.74E-01 |
| BMI | Summed MET minutes/week for all activity | 5.51E-47 | 3.16E-15 | 2.71E-01 |
| BMI | estimated saturated fat intake | 6.44E-03 | 6.37E-01 | 3.85E-01 |
| Diastolic Blood Pressure | Pack years adult smoking as proportion of life span exposed to smoking | 3.03E-04 | 1.75E-04 | 1.08E-02 |
| Pulse Pressure | Pack years adult smoking as proportion of life span exposed to smoking | 7.13E-06 | 8.66E-02 | 4.10E-01 |
| Heart Rate | Pack years adult smoking as proportion of life span exposed to smoking | 3.44E-38 | 5.01E-05 | 3.92E-01 |
| Heart Rate | MET minutes/week for walking | 3.81E-02 | 6.38E-02 | 6.96E-01 |
| Heart Rate | Summed MET minutes/week for all activity | 3.62E-02 | 1.05E-01 | 9.20E-01 |
| Waist-to-Hip Ratio | Pack years adult smoking as proportion of life span exposed to smoking | 5.35E-87 | 6.21E-15 | 7.46E-01 |
| Waist-to-Hip Ratio | MET minutes/week for moderate activity | 9.00E-04 | 2.07E-01 | 2.21E-01 |
| Waist-to-Hip Ratio | MET minutes/week for vigorous activity | 1.13E-04 | 2.46E-01 | 3.12E-01 |
| Waist-to-Hip Ratio | Summed MET minutes/week for all activity | 2.23E-04 | 4.45E-01 | 2.65E-01 |
| Waist-to-Hip Ratio | estimated total energy intake | 2.40E-02 | 1.81E-02 | 1.18E-01 |
| White Blood Cell Count | Pack years adult smoking as proportion of life span exposed to smoking | 1.25E-138 | 3.58E-21 | 6.04E-01 |
| HDL Cholesterol | MET minutes/week for moderate activity | 8.01E-08 | 2.77E-01 | 3.27E-01 |
| HDL Cholesterol | MET minutes/week for vigorous activity | 4.37E-04 | 5.01E-01 | 7.39E-01 |
| HDL Cholesterol | Summed MET minutes/week for all activity | 5.91E-06 | 7.65E-01 | 6.91E-01 |

Only shown for analyses where the full versus null model comparison was significant. Nested models include the null, G-C only and R-C only models. All analyses are based on data after a rank-based inverse normal transformation.

**Sup. Table 7. Genomic relationship within and between top and bottom groups, stratified by G-C interaction estimate, relative to the grand mean genomic relationship.**

| Main Trait | Lifestyle | Within Top Group | | Within Bottom Group | | Between Top & Bottom Groups | |
|---|---|---|---|---|---|---|---|
| | | Δ%[1] | *p-value* [2] | Δ% | *p-value* | Δ% | *p-value* |
| HDL Cholesterol | Physical activity:sports domain | 63.4 | 9.47E-72 | 77.0 | 1.46E-105 | -69.5 | 3.44E-165 |
| HDL3 Cholesterol | Physical activity:sports domain | 64.3 | 2.44E-73 | 67.4 | 3.47E-81 | -66.0 | 2.72E-148 |
| White Blood Cell Count | Physical activity:sports domain | 62.4 | 2.81E-69 | 61.4 | 5.66E-67 | -64.0 | 3.12E-139 |
| HDL2 Cholesterol | Energy from prot1 (%kcal/day) | 84.3 | 2.20E-125 | 87.8 | 4.48E-136 | -82.6 | 4.90E-233 |
| White Blood Cell Count | Physical activity:leisure domain | 67.3 | 5.40E-81 | 66.1 | 2.03E-77 | -65.2 | 4.46E-145 |
| BMI | Saturated fatty acid intake (g/day) | 66.1 | 4.45E-80 | 94.0 | 1.40E-148 | -76.3 | 5.44E-201 |
| HDL3 Cholesterol | Alcohol intake (g/week) | 55.1 | 1.18E-54 | 57.0 | 1.71E-58 | -56.5 | 1.72E-109 |
| Heart Rate | Physical activity:sports domain | 55.2 | 1.72E-54 | 53.2 | 6.04E-52 | -54.1 | 1.72E-101 |

1. Δ% = (group mean - grand mean)/grand mean x 100%; 2. P-values are for two-sided independent t-tests that compare group means with the grand mean

### Supplementary Notes

#### 1. Variance-covariate structure of MRNMs

Following Equation 1 of the text, the main trait, *y*, and the covariate, *c*, for individual *i*, after adjusting for their respective fixed effects, $\mu_y$ and $\mu_c$, are simultaneously expressed as

$$\begin{pmatrix} y_i - \mu_y \\ c_i - \mu_c \end{pmatrix} = \begin{pmatrix} y_i^* \\ c_i^* \end{pmatrix} = \begin{pmatrix} g_i \\ \beta_i \end{pmatrix} + \begin{pmatrix} e_i \\ \varepsilon_i \end{pmatrix}$$

The variance-covariance matrix for *N* realizations of $\begin{pmatrix} y^* \\ c^* \end{pmatrix}$ can be expressed as

$$var\begin{pmatrix} y^* \\ c^* \end{pmatrix} = \begin{bmatrix} \mathbf{Z}_1\mathbf{A}\sigma_{g_1}^2\mathbf{Z}_1' + \mathbf{Z}_1\mathbf{I}\sigma_{e_1}^2\mathbf{Z}_1' & \cdots & \mathbf{Z}_1\mathbf{A}\sigma_{g_{1,N}}\mathbf{Z}_N' + \mathbf{Z}_1\mathbf{I}\sigma_{e_{1,N}}\mathbf{Z}_N' & \mathbf{Z}_1\mathbf{A}\sigma_{g_{1,\beta}}\mathbf{Z}_c' + \mathbf{Z}_1\mathbf{I}\sigma_{e_{1,\varepsilon}}\mathbf{Z}_c' \\ \vdots & \ddots & \vdots & \vdots \\ \mathbf{Z}_\mathrm{N}\mathbf{A}\sigma_{g_{1,N}}\mathbf{Z}_1' + \mathbf{Z}_\mathrm{N}\mathbf{I}\sigma_{e_{1,N}}\mathbf{Z}_1' & \cdots & \mathbf{Z}_N\mathbf{A}\sigma_{g_N}^2\mathbf{Z}_\mathrm{N}' + \mathbf{Z}_N\mathbf{I}\sigma_{e_N}^2\mathbf{Z}_N' & \mathbf{Z}_N\mathbf{A}\sigma_{g_{N,\beta}}\mathbf{Z}_c' + \mathbf{Z}_N\mathbf{I}\sigma_{e_{N,\varepsilon}}\mathbf{Z}_c' \\ \mathbf{Z}_c\mathbf{A}\sigma_{g_{1,\beta}}\mathbf{Z}_1' + \mathbf{Z}_\mathrm{c}\mathbf{I}\sigma_{e_{1,\varepsilon}}\mathbf{Z}_1' & \cdots & \mathbf{Z}_c\mathbf{A}\sigma_{g_{N,\beta}}\mathbf{Z}_N' + \mathbf{Z}_\mathrm{c}\mathbf{I}\sigma_{e_{N,\varepsilon}}\mathbf{Z}_N' & \mathbf{Z}_c\mathbf{A}\sigma_\beta^2\mathbf{Z}_c' + \mathbf{Z}_c\mathbf{I}\sigma_\varepsilon^2\mathbf{Z}_c' \end{bmatrix}$$

Where $\mathbf{I}$ is an *N* x *N* identity matrix, **A** is the *N* x *N* genomic relationship matrix based on genome-wide SNP information, $\mathbf{Z}_\mathrm{i}$ is the incident matrix for $g_i$ for i = 1, 2 …*N*, and $\mathbf{Z}_\mathrm{c}$ is the incident matrix for $\boldsymbol{c}$ .

#### 2. Alternative model comparison strategy

Throughout the text we relied on the null versus full model comparison for detecting G-C and R-C interactions. An alternative and seemingly more logical strategy would be to derive the best model via model comparisons that involve reduced models, i.e., G-C and R-C models, in addition to the null and full models. In this supplementary note we elaborate on this alternative strategy; further show, using results from simulations, issues associated with this strategy, and conclude that the null versus full model comparison is a superior method, hence a logical choice for detecting G-C and R-C interactions.

This alternative strategy uses result patterns from four model comparisons to conclude the best model out of five candidates (Table 1). Candidates considered were the null model, G-C interaction only model, R-C interaction only model, full model, and G-C or R-C interaction models. The last candidate is more of a situation than a model, where the model comparison method does not distinguish between G-C and R-C models, that is, model selection is inconclusive. It occurs when both G-C model and R-C model show a better fit than the null but a worse fit than the full, that is, G-C and R-C models are equally likely. Upon concluding the best model, the source of variance heterogeneity is immediately implied. For example, genetic variance heterogeneity (i.e., a G-C interaction) is declared, when either the G-C model or the full model is the best. In contrast, residual variance heterogeneity (i.e., a R-C interaction) is declared, when either the R-C model or the full model is the best.

**Sup.N2.Table 1. Overview of Model Selection Strategy.**

| Model Comparison | Candidate Model | | | | |
|---|---|---|---|---|---|
| | Null | G-C | R-C | Full | G-C/R-C |
| Null vs. R-C | × | | ✓ | ✓ | ✓ |
| Null vs. G-C | × | ✓ | | ✓ | ✓ |
| R-C vs. Full | | ✓ | × | ✓ | × |
| G-C vs. Full | | × | ✓ | ✓ | × |

Each column shows the model comparison result pattern required to conclude a given candidate model is the best. Result patterns are mutually exclusive across the five candidates. A cross indicates a non-significant p-value for a model comparison (i.e., the simpler model is better), whereas a tick indicates a significant p-value (i.e., the simpler model is worse). Comparisons that are not necessary for model selection are left as blanks. Note the pattern in the last column does not distinguish between G-C and R-C interactions, in which case model selection is inconclusive.

We applied this model selection strategy to simulated data from large- and small-effects settings and evaluated how well the strategy can recover the true simulating model. The results are summarised in Table 2. The method correctly identified the null model for at least 97% of the simulated replicates, giving an estimated type I error rate of 0.03, which is well under the 0.05 target. However, statistical power—estimated by the proportion of replicates for which a true model other than the null is correctly identified—varied largely depending on effect size. For the large-effects setting, a true model other than the null was correctly identified for at least 91% of replicates, hence an estimated power of 0.91 and above. In contrast, for the small-effects setting, the estimated power was 0.04 at worst and 0.11 at best. This does not mean though, the model comparison method could not detect G-C and R-C interactions that are small in magnitude. Rather, for over 75% of replicates under this setting, the likelihood ratio test results were such that G-C and R-C models fit data equally well (see last column of Table 2). In short, either when there are no genuine G-C and R-C interactions or when genuine G-C and R-C interactions are large in magnitude, the likelihood-ratio-based method can discern the true underlying model at a high accuracy (>0.9). However, when genuine G-C and R-C interactions are small, it is unlikely that the method will uncover the true model.

**Sup.N2.Table 2. Proportions of simulated replicates for which a given candidate model is chosen as the best under different simulation scenarios.**

| **Simulation Scenario** | | **Candidate Model** | | | | |
|---|---|---|---|---|---|---|
| | | Null | G-C | R-C | Full | G-C/R-C |
| *large-effects* | No G-C & R-C | **0.98** | 0 | 0 | 0 | 0.02 |
| | G-C only | 0 | **0.93** | 0 | 0.07 | 0 |
| | R-C only | 0 | 0 | **0.96** | 0.04 | 0 |
| | G-C & R-C | 0 | 0.03 | 0.06 | **0.91** | 0 |
| *small-effects* | No G-C & R-C | **0.97** | 0.01 | 0.02 | 0 | 0 |
| | G-C only | 0.06 | **0.11** | 0.03 | 0.04 | 0.76 |
| | R-C only | 0.1 | 0.01 | **0.09** | 0.05 | 0.75 |
| | G-C & R-C | 0 | 0.05 | 0.14 | **0.04** | 0.77 |

Data were simulated under normality with large and small effects in terms of heritability, G-C and R-C interactions. Each scenario had 100 replicates of a main trait and a covariate. For each replicate, four models were fitted and compared to select the best fitting one (see Table 1 of this Supplementary Notes).

For each simulation scenario, we also compared parameter estimates from all four fitted models with their corresponding true values and noted that results are similar for the two parameter settings that vary in effect sizes. Figure 1 in this supplementary note shows results for the small-effects setting, which hold for the large-effects setting. When the true underlying model was the null, regardless of which model was fitted, all parameter estimates were unbiased. In scenarios where the true model was other than the null, fitting the correct model produced unbiased estimates for all parameters. However, in these scenarios, fitting a wrong model—that is, a model other than the true—could produce biased estimates for some parameters. For example, fitting the null model to data with genuine G-C or/and R-C interactions produced larger estimates of the residual variance, i.e., $\delta^2_{\tau 0}$, than its true value, by an amount similar to the set value of $\delta^2_{\alpha 1}$ or/and $\delta^2_{\tau 1}$. When fitting the G-C model to data with R-C interaction but no G-C interaction, estimates of $\delta^2_{\alpha 1}$ were larger than the true, i.e., 0, by an amount similar to the set value of $\delta^2_{\tau 1}$. Likewise, when fitting the R-C model to data with G-C interaction but no R-C interaction, estimates of $\delta^2_{\tau 1}$ deviated from the true by an amount similar to the set value of $\delta^2_{\alpha 1}$. However, fitting the full model, even when it was the wrong model, produced unbiased estimates for all parameters. Thus, our simulation results indicate that model misspecification can result in biased estimates for some parameters depending on the simulation scenario, with the only exception of fitting the full model, which provides unbiased estimates for all parameters in all scenarios.

In summary, the model selection strategy has a type I error rate under 0.05, but its power of recovering the true model is very low when effect sizes are small. Consequently, this strategy can result in an alarmingly elevated chance of concluding a wrong model, i.e., model misspecification, which could produce biased estimates for some model parameters. Therefore, for analysis of real data, where the true underlying model is unknown and effects sizes are likely small, this model comparison strategy is not useful to select the best model for identifying source of variance heterogeneity. In contrast, we showed in the main text that the null versus full model comparison method has an acceptable type I error rate and reasonable power when effect sizes are small. Even if the full model is not true, model estimates are not biased, which can be interpreted subsequently. Hence, the null versus full model comparison is a superior method to the alternative model comparison strategy.

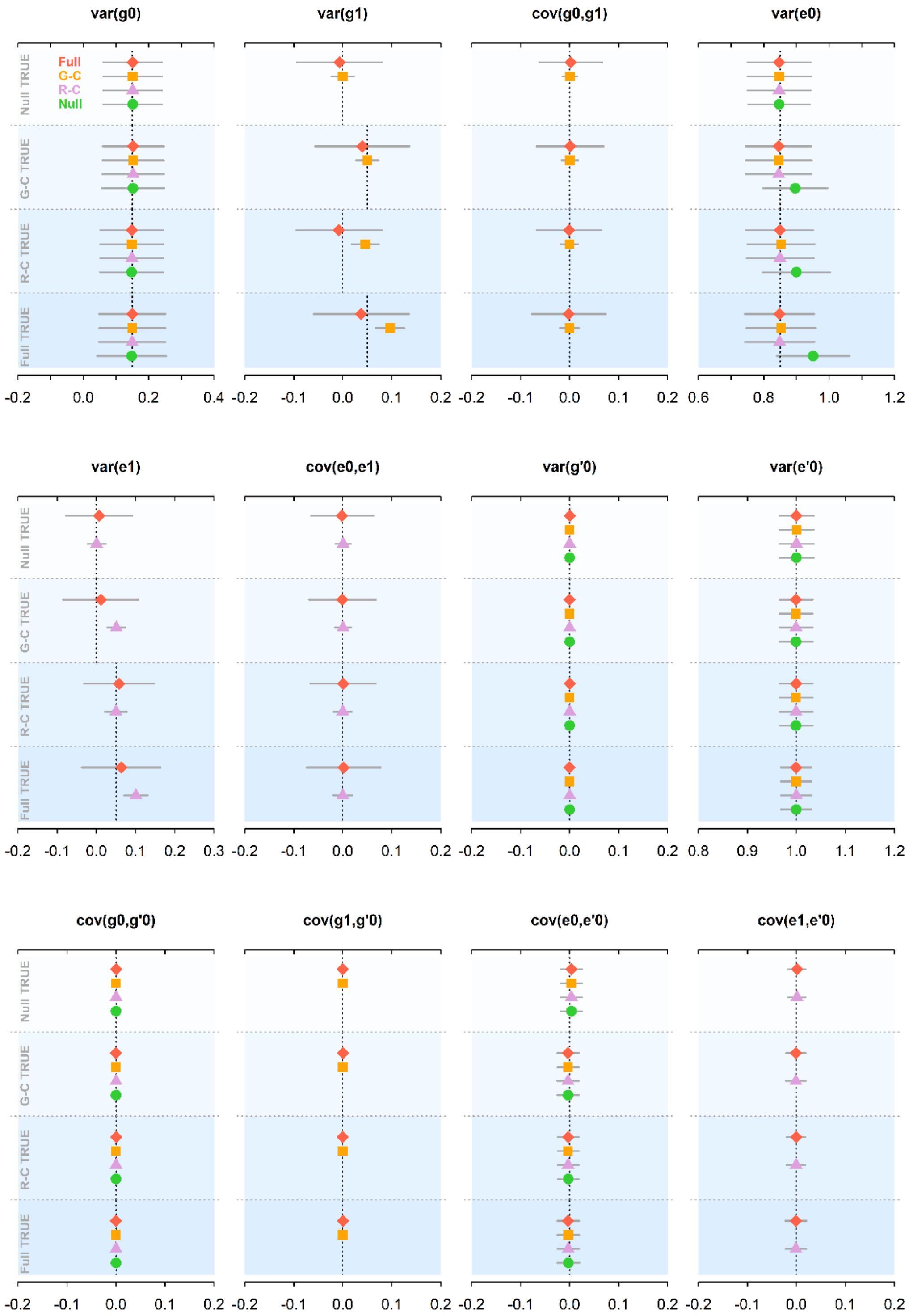


**Sup.N2.Figure 1. Sampling distributions of parameter estimates for four multivariate reaction norm models under four simulation scenarios for the small-effects parameter setting.** The four scenarios are no G-C and R-C interactions (i.e., a null model), G-C interaction only (i.e., a G-C model), R-C interaction only (i.e., a R-C model), and both G-C and R-C interactions (i.e., a full model). There are 100 replicates under each scenario, with the true value of each parameter being indicated by a vertical dashed line.
